# An event-based paradigm for analyzing fluorescent astrocyte activity uncovers novel single-cell and population-level physiology

**DOI:** 10.1101/504217

**Authors:** Yizhi Wang, Nicole V. DelRosso, Trisha Vaidyanathan, Michael Reitman, Michelle K. Cahill, Xuelong Mi, Guoqiang Yu, Kira E. Poskanzer

## Abstract

Recent work examining astrocytic physiology centers on fluorescence imaging approaches, due to development of sensitive fluorescent indicators and observation of spatiotemporally complex calcium and glutamate activity. However, the field remains hindered in fully characterizing these dynamics, both within single cells and at the population-level, because of the insufficiency of current region-of-interest-based approaches to describe activity that is often spatially unfixed, size-varying, and propagative. Here, we present a paradigm-shifting analytical framework that releases astrocyte biologists from ROI-based tools. Astrocyte Quantitative Analysis (AQuA) software enables users to take an event-based approach to accurately capture and quantify the irregular activity observed in astrocyte imaging datasets. We apply AQuA to a range of *ex vivo* and *in vivo* imaging data, and uncover previously undescribed physiological phenomena in each. Since AQuA is data-driven and based on machine learning principles, it can be applied across model organisms, fluorescent indicators, experimental modes, and imaging resolutions and speeds, enabling researchers to elucidate fundamental astrocyte physiology.

## Introduction

With increased prevalence of multiphoton imaging and optical probes to study the physiology of astrocytes^1–3^, many groups now have the tools to study fundamental functions that previously remained unclear. Recent work has focused on new ways to decipher how astrocytes respond to neurotransmitter and neuromodulator circuit signals^4–7^ and how the spatiotemporal patterns of their activity shape local neuronal activity^8–10^. Recording astrocytic dynamics with the goal of decoding their disparate roles in neural circuitry has largely centered on cell type-specific expression of genetically encoded probes to carry out calcium (Ca^2+^) imaging using variants of GCaMP^3^, and glutamate imaging using GluSnFR^2^. Compared to neuronal Ca^2+^ imaging, astrocytic Ca^2+^ imaging using GCaMP presents particular challenges due to their complex spatiotemporal dynamics. Thus, astrocyte-specific analysis software has been developed to capture these dynamics, including techniques that divide the cell into distinct subcellular regions corresponding to their anatomy^4^ or apply a watershed algorithm to identify regions-of-interest (ROIs)^11^. Likewise, GluSnFR imaging analysis techniques are based on manually or semi-manually selected ROIs, or by analyzing the entire imaging field together as one ROI. It is worth noting that these and other current techniques rely on the conceptual framework of ROIs for image analysis. However, astrocytic Ca^2+^ and GluSnFR fluorescence dynamics are particularly ill-suited for ROI-based approaches, because the concept of the ROI has several inherent assumptions that cannot be satisfied for astrocytic activity data. Astrocytic Ca^2+^ signals, for example, can occupy regions that change size or location across time, can propagate within or across cells, and can spatially overlap with other Ca^2+^ signals that are temporally distinct. ROI-based approaches assume that for a given ROI, all signals have a fixed size and shape, and all locations within the ROI undergo the same dynamics, without propagation. Accordingly, ROI-based techniques may over- or under-sample these data, thus obscuring true dynamics and hindering physiological discovery in these cells. An ideal imaging analysis framework for astrocytes would take into account, and quantify, all of these dynamic features and be free of these ROI-based analytical restrictions. In addition, the ideal tool would be applicable to astrocyte imaging data across spatial scales, encompassing subcellular, cellular, and population-wide fluorescence dynamics.

In this work, we set out to design an image analysis toolbox that would capture the complex, wide-ranging fluorescent signals observed in most dynamic astrocyte imaging datasets. We reasoned that a non-ROI-based approach would best describe the observed fluorescent dynamics, and applied probability theory, machine learning, and computational optimization techniques to generate an algorithm to do so. We name this resulting software package Astrocyte Quantitative Analysis (AQuA) and validate its utility by applying it to simulated datasets that reflect the specific features that make analyzing astrocyte data challenging. We next apply AQuA to three experimental two-photon (2P) imaging datasets*—ex vivo* Ca^2+^ imaging of GCaMP6 from acute cortical slices, *in vivo* Ca^2+^ imaging of GCaMP6 in primary visual cortex (V1) of awake, head-fixed mice, and *ex vivo* glutamate imaging of both astrocytic and neuronal expression of GluSnFR. In these test cases, we find that AQuA accurately detects fluorescence dynamics by capturing fluorescence events as they change in space and time, rather than the activity from a single location in space, as in ROI-based approaches. AQuA outputs a comprehensive set of biologically relevant parameters from these datasets, including propagation speed, propagation direction, area, shape, and spatial frequency. Using these detected events and associated output features, we uncover neurobiological phenomena that have not been previously described in astrocytes. A wide variety of cellular and circuit functions have been ascribed to astrocytes, and a key question currently under examination in the field is whether certain classes of Ca^2+^ or glutamate activities observed in these cells correspond to particular neurobiological functions. The framework we describe here allows for a rigorous, in-depth dissection of astrocyte physiology across spatial and temporal imaging scales, and sets the stage for a comprehensive categorization of heterogeneous astrocyte activities both at baseline and after experimental manipulations.

## Results

### Design principles of the AQuA algorithm

To move away from ROI-based analysis approaches and accurately capture heterogeneous astrocyte fluorescence dynamics, we designed an algorithm to decompose raw dynamic astrocyte imaging data into a set of quantifiable events (Supp. Fig. 1–2). An event is a signal transient occurring in a certain region, but this region is defined by the fluorescent dynamics, not *a priori* by the user or the cell morphology. Since an event is defined by transient changes in fluorescence, a location in the imaging field can be associated with multiple events over the course of the imaging period, as the same location may undergo multiple fluorescence transients. Importantly, this definition means that events are not necessarily spatially fixed; they may occupy different regions over time. Methodologically, ROI-based analyses restrict multiple transients to the same region. We argue that this restriction is too strong and thus not matched to dynamic astrocyte fluorescence data, as evidenced by the well documented observation that Ca^2+^ activity may involve the whole astrocyte (including soma and all processes), or may be constrained to a small segment of a process^6,8,12^. Here, we directly model these unfixed events in the algorithm, which omits the concept of ROI entirely. Our event-based approach not only resolves difficulties inherent in ROI-based analyses, but it also provides richer information with which to characterize relevant astrocytic physiology since the dynamic regions associated with different events are quantifiable.

We mathematically define an event as a cycle of a signal increase and decrease that coherently occurs in a spatially connected region, and therefore is specified jointly by its spatial and temporal properties. Importantly, our definition can flexibly accommodate the common phenomenon of fluorescence propagation in astrocyte imaging data, because transients at different locations are required to be coherent but not necessarily synchronized. In our algorithm, an event must satisfy the following two rules: 1) the temporal trajectory for an event contains only one peak (single-cycle rule, Fig. 1a) and 2) adjacent locations in the same event have similar trajectories (smoothness rule, Fig. 1a). Briefly, our strategy of event-detection is to a) explore the single-cycle rule to find peaks, which are used to specify the time window and temporal trajectory, b) explore the smoothness rule to group spatially adjacent peaks, whose locations specify the occupied region, c) apply machine learning and optimization techniques to iteratively refine the spatial and temporal properties of the event to best fit the data, and d) apply statistical theory to determine whether a detected event is true or due to noise (Fig. 1). Full statistical and computational details are provided in the Methods, but we want to highlight one technical innovation and one new concept that jointly enable a nuanced analysis of astrocyte fluorescence dynamics as shown below in application to experimental datasets. The technical innovation is our development of the mathematical model Graphical Time Warping^13^ (GTW), with which we are able to consider fluorescent signal propagation as integrated into each modeled event. To the best of our knowledge, signal propagation has never been rigorously accounted for and has been considered an obstacle to analysis. With GTW, we can estimate and quantify propagation patterns in the data. In addition, we introduce a third rule (single-source rule (Fig. 1a)) such that each event only contains a single initiation source. With the single-source rule, we can separate events that are initiated at different locations but meet in the middle. The single-source rule also allows us to divide large bursts that occur across a field-of-view into individual events, each with a single initiation location.

**Figure 1.**
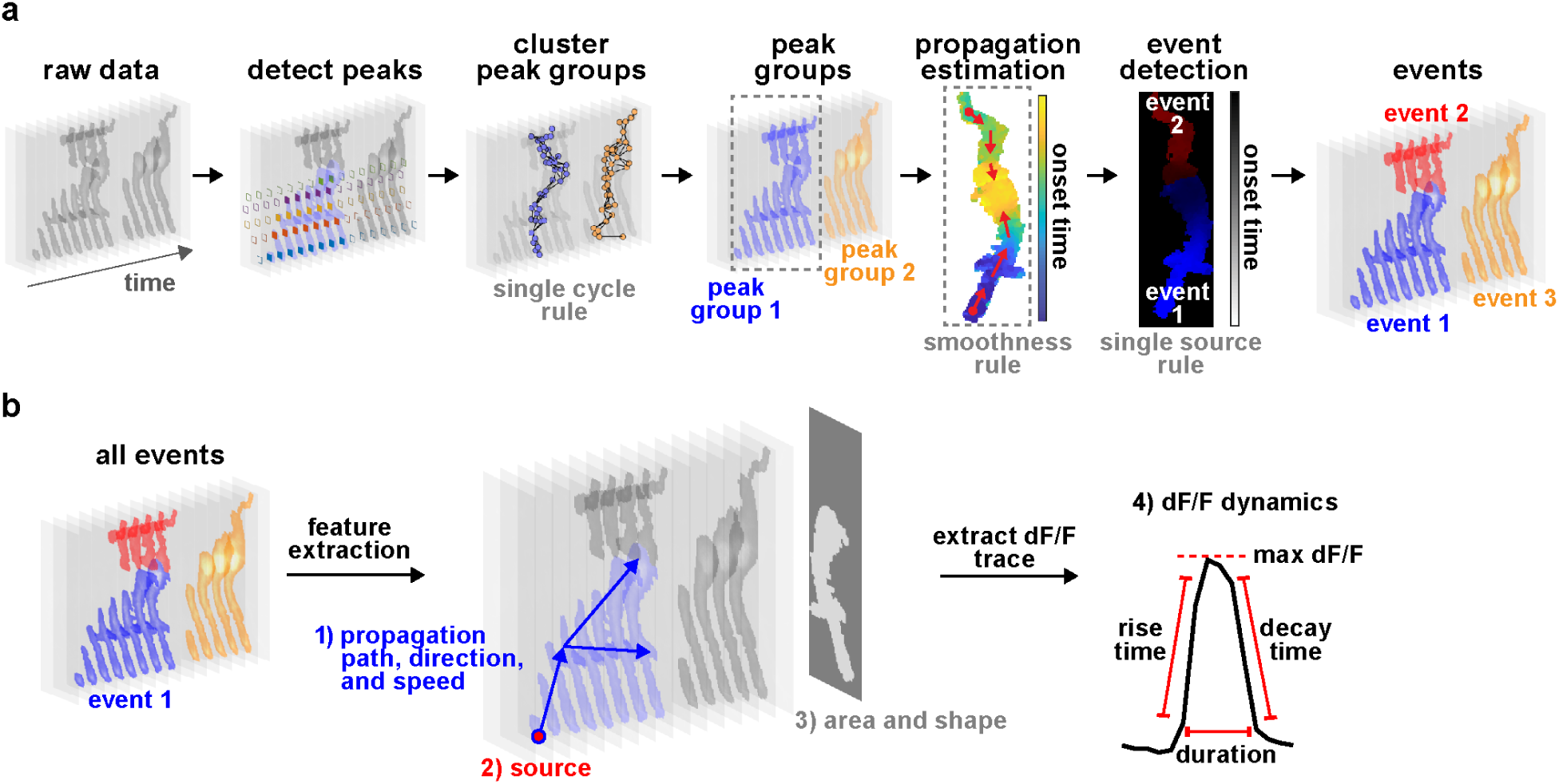
AQuA algorithm. **(a)** Flowchart of AQuA algorithm. Raw data is visualized as a stack of images across time with grey level indicating signal intensity. In the *detect peaks* panel, five peaks are detected and highlighted by solid diamonds, each color denoting one peak. Based on the single-cycle rule and spatial adjacency of the apexes of each peak, peaks are clustered into spatially disconnected groups. Apexes are labelled as solid dots. Based on smoothness, propagation patterns are estimated for each peak group. By applying the single-source rule, two events are detected for peak group 1. Three total events are detected. **(b)** Feature extraction. Based on the event-detection results, AQuA outputs four sets of features relevant to astrocytic activity: 1) propagation related (path, direction, and speed); 2) source of events, indicating where an event is initiated; 3) features related to the event footprint, including area and shape. Event 2 is plotted here; 4) features derived from the dF/F dynamics.

**Figure 2.**
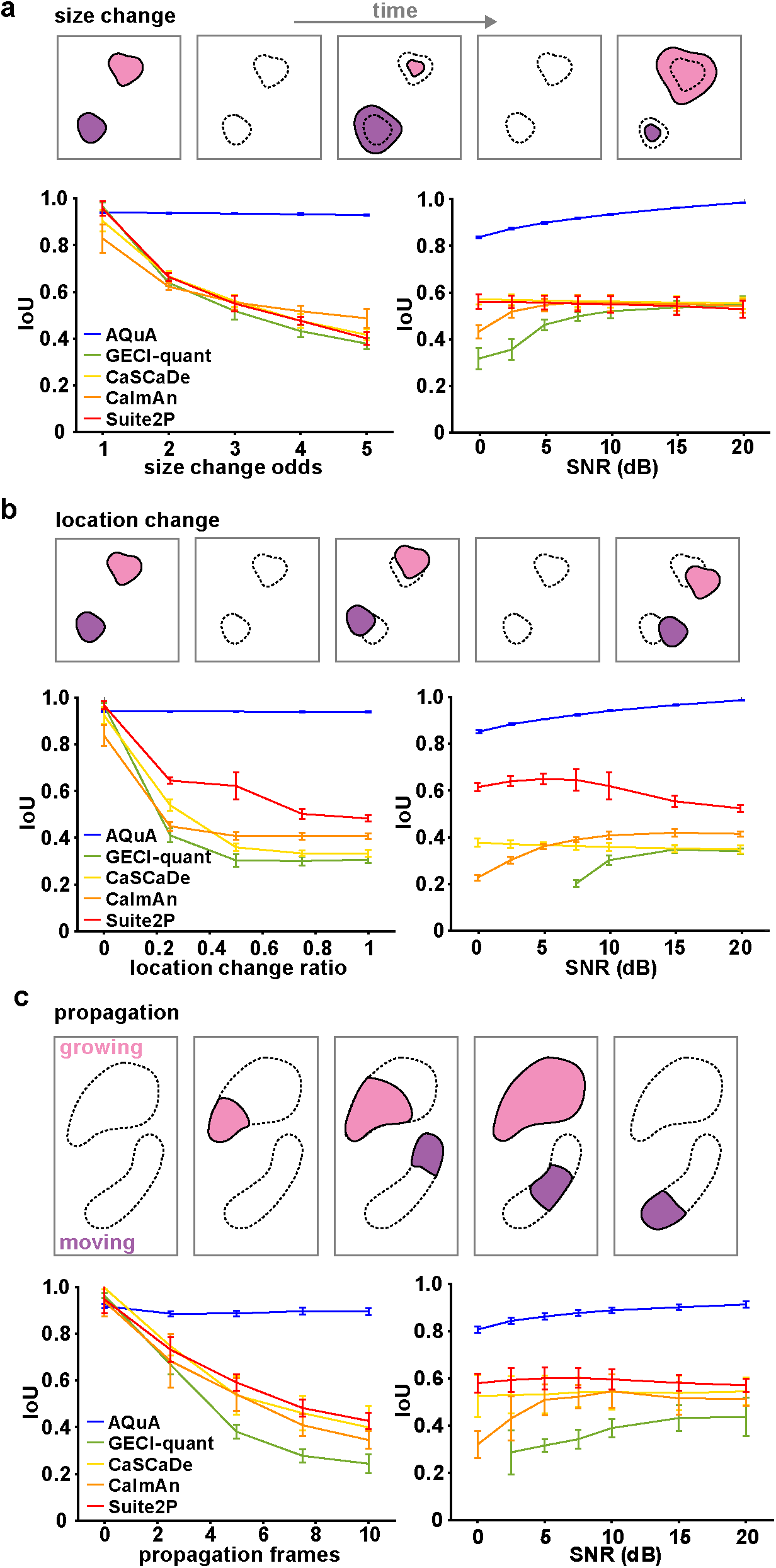
Performance comparison among image-analysis methods. **(a–c)** Schematic (top) and results (bottom) of performance of five image analysis methods (AQuA, GECI-quant, CaSCaDe, CaImAn, and Suite2P) on simulated datasets, independently changing event size (a), location (b), and propagation duration (c). In results, change of independent parameter is shown in left panel, and varying SNR in right. For each result, the smallest value of the independent parameter corresponds to a simulation under pure ROI assumptions. The larger the values, the greater the violation of the ROI assumptions. IoU (intersection over union) measures the overlap between detected and ground-truth events. An IoU=1 is the best achievable performance, meaning that all detected events are ground-truth and all ground-truth events are detected. Error bars indicate the 95% confidence interval calculated from 10 independent replications of simulation, where each simulation contains hundreds of events.

The output of the AQuA algorithm is a list of detected events, each associated with three categories of parameters: 1) the spatial map indicating where the event occurs, 2) the dynamic curve corresponding to fluorescence change over time (dF/F), and 3) the propagation map indicating signal propagation. For each event, we use the spatial map to compute the event area, diameter and shape of the domain it occupies (Fig 1b). Using the dynamic curve, we can calculate maximum Δ*F/F*, duration, onset-time, rise-time and decay-time. Using the propagation map, we extract event initiation location, as well as propagation path, direction, and speed. A complete list of features is in the Methods section.

### Validation of AQuA using simulated data

To validate AQuA, we designed three simulation datasets for which we have complete knowledge of when, where, and how each event occurred. These three datasets correspond to the three key ROI-approach-incompatible phenomena: size-variability, location-variability, and propagation. While these three phenomena usually co-occur in real datasets, we simulated each phenomenon independently to examine their individual impact and test AQuA’s performance relative to other fluorescence image analysis tools, including CaImAn^14^, Suite2P^15^, CaSCaDe^11^, and GECI-quant^4^. CaImAn and Suite2P are widely used for neuronal Ca^2+^ imaging analysis while CaSCaDe and GECI-quant were designed specifically for Ca^2+^ activity in astrocytes. Although CaSCaDe can output events, all four methods are ROI-based.

We first studied the impact of size-varying events (Fig. 2a), in which multiple events occur at the same location and the event centers remain fixed, but event sizes change. The degree of size change is quantified using size-change odds (see Methods) where a size-change odds of 1 indicates events with the same size, while an odds of 5 is the largest size change we simulated. When we set the odds at 5, we simulate events with sizes randomly distributed between 0.2 and 5 times the baseline size, with an SNR of 10dB, chosen to closely match the noise level in real experimental data. Two measures, IoU and event count, were then used to evaluate the performance on all simulated datasets. IoU (intersection over union) indicates the overlap between detected and the ground-truth events, and takes into account both the spatial and temporal accuracy of detected events. An IoU of 1 is the best performance possible, while an IoU of 0 is the worst. When there is no size change (odds=1), all methods have good IoU performance around 0.95, with CaSCaDe and CaImAn slightly worse than others (Fig. 2a). When the degree of size change is increased, AQuA still performs well (IoU=0.95), while all other methods quickly drop to 0.4–0.5. We then changed our analysis to study the impact of different SNRs on performance by varying SNR, but fixing the size-change odds. AQuA performed better with increasing SNR and achieved nearly perfect detection accuracy (IoU=1) at 20dB. In comparison, all other methods had an IoU below 0.6, even at high SNR (Fig. 2a). We also examined the results by visualizing event counts for each pixel (Supp. Fig. 3–4). With size change odds of 3 or 5, the map of ground truth event-counts did not show clear ROI boundaries, because events from the same ROI had various sizes, and because events from different ROIs can overlap at some spatial locations (Supp. Fig. 3). It is clear from these maps that AQuA reported faithfully the events under various SNRs but all other methods had erroneous event counts and produced artificial patterns. The visualization also informed us different types of errors in other methods. CaSCaDe tends to over-segment, as it is based on watershed segmentation. GECI-quant, especially its soma-segmentation step, is particularly challenged by noise, causing many signals to be lost (Supp. Fig. 4).

We next focused on the impact of shifting the event locations. In these simulated datasets, event size is fixed but event location changes, and degree of change is represented by a location change ratio (Fig. 2b). A ratio of zero indicates no location change. Here, results are similar to changing size, as above. AQuA modeled the location change well and its performance was not affected by degree of location change. Likewise, AQuA reached near perfect results when SNR was high. In contrast, all other analysis methods performed poorly with changing locations. In particular, the other astrocyte-specific methods (CaSCaDe and GECI-quant) missed many signals. Even though the overall conclusion is similar for both the size- and location-changing events, the peer methods had more variation of IoU performance among themselves and the event count map showed distinct patterns (Supp. Fig. 3). In general, changing event locations is more challenging for ROI-based methods because when the location change is large, two events may have no spatial overlap, which is never true for size-varying events. This accounts for results seen when applying GECI-quant for example, including the result that GECI-quant is not able to detect anything when the SNR is low (Fig. 2b, right).

In our third simulated dataset, we asked how the phenomenon of fluorescence signal propagation impacts the performance of AQuA compared to the other methods. Two propagation types—growing and moving—were simulated in this dataset (Fig. 2c), although they were also separately evaluated (Supp. Fig 5). Propagation frame number denotes the difference between the earliest and latest onset times within a single event. When propagation frame number is zero, all signals within one ROI, but not necessarily across ROIs, are synchronized and there is no propagation. Similar results to the two scenarios discussed above were obtained here, with AQuA out-performing all the other methods by a large margin. These results indicate that AQuA can handle various types of propagation well, while the performance of other methods degrades rapidly when propagation is introduced. We note that although all events are constrained in the same ROI, propagation caused ROI-based approaches to quickly decline in performance. GECI-quant was again influenced by noise level, while CaSCaDe’s assumption of synchronized signals did not allow accurate capture of the event dynamics.

In all, when any of the three ROI-violating factors—size-variability, location-variability, and propagation—is introduced, other methods do not accurately capture the signal dynamics, and AQuA outperforms them by a large margin. We expect that the performance margins on experimental data is larger than those quantified in the simulation studies here, since real data exhibits multiple ROI-violating factors and the performance of the ROI-based methods is over-estimated in our simulations (see Methods).

### AQuA enables identification of single-cell physiological heterogeneities

To test AQuA’s performance on real astrocyte fluorescence imaging data, we first ran AQuA’s event-detection on Ca^2+^ activity recorded from astrocytes in acute cortical slices from mouse V1 using 2P microscopy. We used a viral approach to express the genetically encoded Ca^2+^ indicator GCaMP6f^3^ in layer 2/3 (L2/3) astrocytes. Unlike ROI-based approaches that use arbitrarily sized shapes to extract fluorescence, AQuA detects both propagative and non-propagative activity, revealing Ca^2+^ events with a variety of shapes and sizes (Fig. 3a, left). Further, since AQuA not only detects Ca^2+^ events’ spatial footprint but also their time-course, we can apply AQuA to measure the propagation direction each event travels over its lifetime. Imaging single cells, we used the soma as a landmark, and classified events as traveling toward the soma (pink), away from the soma (purple), or static (blue) for the majority of its lifetime (Fig. 3a, right). We used AQuA’s automatic feature-extraction and combined multiple measurements (size, propagation direction, duration, and minimum proximity to soma) into one spatiotemporal summary plot (Fig. 3b). Since astrocytes exhibit a wide diversity of Ca^2+^ activities across subcellular compartments^6,16,17^, plotting the signals this way rather than standard dF/F transients highlights these heterogeneities, allows us to map the spatial location of the Ca^2+^ signals, and enables a quick, visual impression of a large amount of complex data (Supp. Fig. 6).

**Figure 3.**
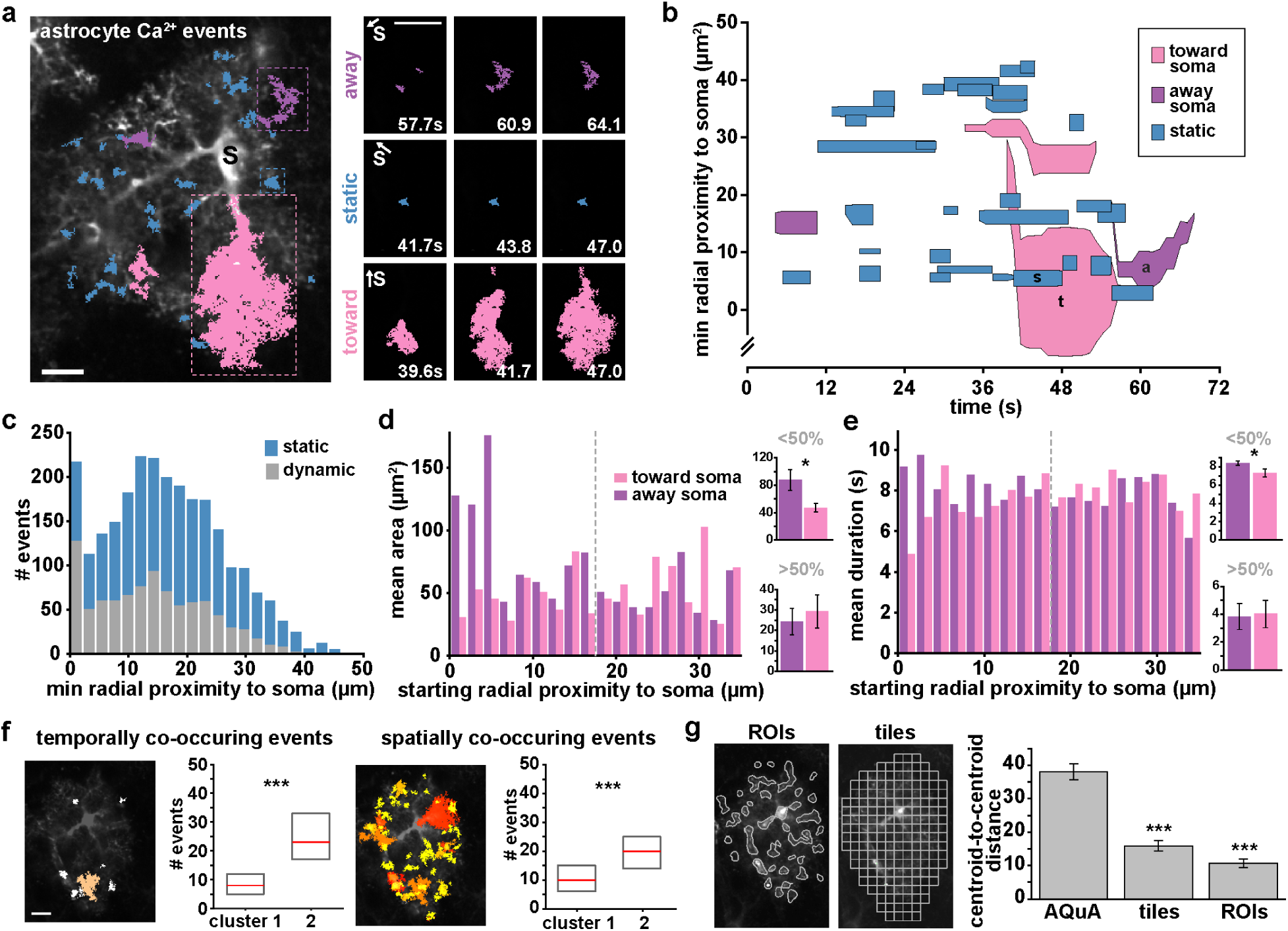
AQuA features capture heterogeneities among single astrocytes. **(a)** Representative GCaMP6f *ex vivo* image (left) with AQuA events overlaid from 1 min of a 5 min movie. Soma marked with black s. (Scale bar = 50μm; Supplemental movie 1). Right: Representative image sequence for each propagation direction class (blue=static, pink=toward soma, purple=away from soma; scale bar=20μm. Soma direction marked with s and white arrow. **(b)** Spatiotemporal plot of Ca^2+^ activity from 1 min of movie. Each event is represented by a polygon that is proportional to its area as it changes over its lifetime. **(c)** Distribution of dynamic and static events as a function of minimum distance from soma (chi-square test, ***p<0.001, n=5 slices, 11 cells). All bin widths calculated by Freedman-Diaconis’s rule. **(d)** Left: Propagative event size versus starting distance from soma, segregated by propagation direction. Dashed gray line denotes half the distance between the soma and the cell border. Right: Average event area for those that start <50% (top) and >50% (bottom) of the distance from the soma, (one-tailed paired t-test, *p<0.05). **(e)** Left: Event duration versus starting distance from soma. Right: Average event duration for those that start <50% (top) and >50% (bottom) of the distance from the soma (one-tailed paired t-test, *p<0.05). **(f)** Two event-based measurements of frequency: events with activity overlapping in time (top) and in space (bottom; scale bar=50μm). Median (red) and interquartile range (blue) from the three cells in cluster 1 and the eight cells in cluster 2 (one-tailed Wilcoxon rank sum, ***p<0.001). **(g)** After t-SNE plotting of Ca^2+^ activity using features calculated from ROIs and 5×5μm tiles (top), quantification of centroid distances between cells from cluster 1 and cluster 2 (bottom, one-tailed paired t-test, ***p<0.001).

We next asked whether some subcellular regions of astrocytes have more dynamic activity than others across all analyzed cells (n=11 cells). Although we detected more static events than dynamic overall (Supp. Fig. 7a), we observed a higher proportion of dynamic events than static events in the soma (59%, Fig. 3c, Supp. Fig. 7b). We then characterized events by propagation direction and event initiation location (Fig. 3d). Events that begin close to the soma (≤50th percentile) and propagate away (purple) were on average larger than the events propagating toward the soma (pink, two-tailed t-test). Similarly, those events that began close to the soma and propagated away had on average a longer duration than events propagating toward the soma (two-tailed t-test, Fig. 3e, Supp. Fig. 7).

One of AQuA’s strengths is its ability to automatically extract large numbers of features. These features can be used to form a comprehensive Ca^2+^ measurement matrix. Dimensionality reduction applied to this matrix can, in turn, be used to visualize each cell’s Ca^2+^ signature (Supp. Fig. 8). To do this, we applied t-distributed Stochastic Neighbor Embedding (t-SNE)^18^, followed by k-means clustering to assign the cells to groups (Supp. Fig. 8), revealing two clusters marked by cells with large differences in median frequency (Fig. 3f). Astrocytic Ca^2+^ frequency is commonly measured as the number of transients that occur over time within an ROI. Here, we instead define frequency from an event-based perspective in two ways: 1) for each event, the number of other events that overlap in time, and 2) for each event, the number of other events that overlap in space. We used these two measures (temporal and spatial overlap) and several other extracted measures (Supp. Fig. 8) to construct the matrix used for t-SNE visualization and clustering. We next tested how well our AQuA-specific features perform at clustering the heterogeneity between cells compared to two ROI-based methods (Fig. 3g), and found that the AQuA-based method outperformed the others. In fact, even when we only use AQuA-specific features for this analysis—area, temporal overlap, spatial overlap, and propagation speed—and remove all features that can be extracted from ROI-based methods, AQuA still significantly outperforms in clustering cells (Supp. Fig. 8g–i)). AQuA-extracted features that correspond only to those that can be obtained by ROI-based methods—frequency, amplitude, duration—do not allow clustering significantly better than the ROI-based approaches themselves (Supp. Fig. 8g–i), suggesting that the AQuA-specific features are those that best capture dynamic fluorescence features that vary among single cells. This indicates that AQuA can be used to extract data from existing *ex vivo* Ca^2+^ imaging datasets to reveal previously uncovered dynamics and sort cells into functionally relevant clusters.

### In vivo astrocytic Ca^2+^ bursts display anatomical directionality

Recent interest in astrocytic activity at the mesoscale has been driven by population-level, multi-cellular astrocytic Ca^2+^ imaging^1,5,7,8,12,19,20^. To test the power of AQuA-based event detection, we next applied it to populations of *in vivo* astrocyte Ca^2+^ activity. Previous studies have described temporal details of astrocyte activation^4,5,7,8,12^, yet have left largely unaddressed the combined spatiotemporal properties of Ca^2+^ activity at the circuit-level. Here, we explored whether AQuA can uncover spatial patterns within populations of cortical astrocytes in an awake animal, and carried out head-fixed, 2P imaging of GCaMP6f activity in V1, L2/3 astrocytes. To minimize motion artifacts, we first registered our imaging sets using non-rigid motion correction^21^. Populations of *in vivo cortical* astrocytes exhibit both small, focal, desynchronized Ca^2+^ activity, and large, synchronous activities^4,5^. AQuA detected both of these classes of Ca^2+^ activity (Movie 2, Fig. 4a). Similar to previous studies, we observed many (but not all) of the synchronous bursts co-occurring with locomotion periods (Fig. 4b, pink), and many events within these burst periods displayed propagation (Fig. 4c, top). These propagative events were larger in area and had greater propagation distances compared to the events that occurred during the inter-burst periods (Fig. 4c, bottom). Here, to examine a distinct type of neurobiological phenomenon and test whether AQuA could help us analyze discrete features of this phenomenon, we focused our investigation only on these events occurring during the burst periods (Supp. Fig. 9).

**Figure 4.**
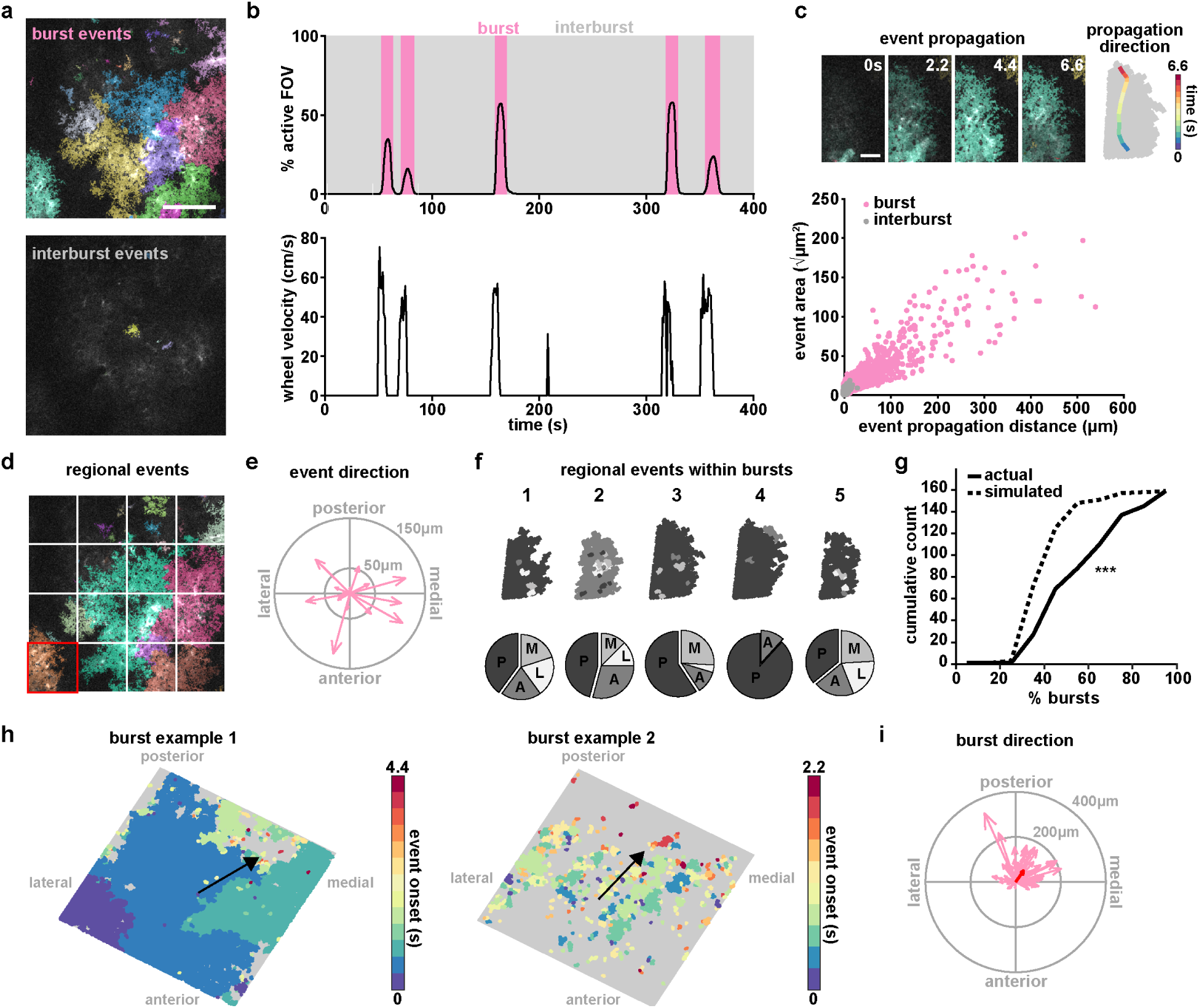
AQuA resolves astrocytic Ca^2+^ propagation directionality across scales. **(a)** Representative *in vivo* GCaMP6f images during a burst period (top) and inter-burst period (bottom) with overlaid AQuA-detected events (scale bar=50μm). **(b)** Population Ca^2+^ events represented as percentage of the imaging field active as a function of time. Burst periods (pink) are identified when Ca^2+^ activity exceeds more than 1% of the active field of view and exceeds more than 10% of the maximum number of event onsets. **(c)** *In vivo* Ca^2+^ events propagate with specific directionality. Top: representative propagative event that occurred during the burst period in panel a. (scale bar=25μm). The propagation direction (change of centroid relative to its original location) for each frame is overlaid on the event (right). Bottom: Total propagation distance versus event size for all events within bursts (n=6 mice, 66 bursts, 14,967 events). **(d)** To test consistency of local directionality during bursts, sixteen 96×96μm tiles are overlaid on images. **(e)** Event propagation direction from all events over the entire field in the burst shown in *d*. Length of arrow indicates propagation distance. **(f)** Top: All events within highlighted tile in *d* (*red square*) for five burst periods, color-coded by propagation direction (top). Bottom: Event propagation direction distributions (P=posterior; A=anterior; M=medial; L=lateral). **(g)** Cumulative distribution of percentage of bursts with events (within individual tiles/regions) propagating in the same direction in actual (solid) and simulated (dashed) data (one-tailed Wilcoxon rank sum, ***p<0.001) **(h)** Two representative maps of population burst propagation direction with each event color-coded by their onset time relative to the beginning of the burst period, demonstrating variability of burst size. **(i)** Burst propagation direction calculated from onset maps in *h* (n=66 bursts). Event locations from the first 20% of the frames after burst onset are averaged together to determine burst origin. Event locations from 20% of the last frames after burst onset are averaged together and the difference between this and the origin determines burst propagation distance. Red arrow denotes average of all bursts.

To analyze burst-period Ca^2+^ events, we first interrogated the consistency of event propagation direction across all burst periods. To do so, we divided our field-of-view into 16 equivalently sized, regional tiles (Fig. 4d). Within a single burst, plotting individual event direction within the entire field of view did not reveal a consistent propagation direction (Fig. 4e). However, within a single region, the major propagation direction across bursts was consistent (Fig. 4f). When we plot the cumulative count of the percentage of bursts with regions that propagate in the same direction, we indeed observe that this curve is right-shifted compared to a simulated random assignment of majority regional propagation direction (Fig. 4g). Thus, local, small-scale fluorescence activity exhibits directional consistency across bursts.

Because the percentage of the active field of view varied across burst periods (Fig. 4b), with a wide variability from few to hundreds of events (Fig. 4h), we also measured the propagation direction of the events within each burst period, now using each event’s onset time to calculate a single burst-wide propagation direction (Fig. 4h, black arrow). Doing so revealed a consistent posterior-medial directionality of population Ca^2+^ activity in L2/3 V1 astrocytes (Fig. 4i). Although Ca^2+^ bursts have been previously observed using GCaMP6 imaging in awake mice^4,5^, consistent spatial directionality with respect to the underlying anatomy has never been described. This observed posterior-medial directionality may be revealing anatomical and physiological underpinnings of these bursts, and since they have been shown to be at least partly mediated by norepinephrine^5,7^, they could be reflective of the response of groups of cortical astrocytes to incoming adrenergic axons originating in locus coeruleus.

### Astrocytic and neuronal expression of GluSnFR reveals differential glutamate dynamics

We next asked whether AQuA could be used to detect astrocytic fluorescent activities with very different spatiotemporal dynamics than we observe when measuring intracellular Ca^2+^. We decided to carry out GluSnFR imaging^22^ to measure extracellular glutamate dynamics, since GluSnFR has been widely used for glutamate imaging^2,6,8^ and one canonical function of astrocytes is to regulate extracellular glutamate. While GluSnFR has been expressed both in astrocytes and in neurons previously^2,20,21,2,8,22,23^, how cell type-specific expression determines its fluorescent dynamics has not been fully explored. No previously applied analytical tools have been reported to automatically detect GluSnFR-based glutamate events to accommodate differential event sizes and shapes. Here, we explored whether application of AQuA could be used to detect cell type-specific differences in glutamate dynamics and help reveal heterogeneities of glutamate events based on various underlying biological mechanisms.

We expressed GluSnFR in either astrocytes or neurons using injections of cell type-specific viruses (*AAV1-GFA*(*ABC*(*1*)*D*)*-iGluSnFR* and *AAV1-hsyn-iGluSnFR*, respectively) and carried out 2P imaging of spontaneous activity in acute cortical slices. Distinct morphological differences between astrocytic and neuronal expression of GluSnFR were evident, as has been observed previously^8,24,25^ (Fig. 5a). We next applied AQuA to these datasets to detect significant increases in GluSnFR fluorescence, and were able to detect events that were too small and dim to detect by eye (Movie 3). Indeed, 62% of astrocytic events (n=157) had an area less than the size of a single astrocyte (100 μm^2^), and 8% of astrocytic (n=157) and 35% of neuronal glutamate events (n=107) had a maximum dF/F less than 0.5. Because GluSnFR events have previously been detected by spatially averaging within a single cell or across broader areas of tissue, *or* by manual detection, the events that AQuA detects are missed by these other methods (Supp. Fig. 10).

**Figure 5.**
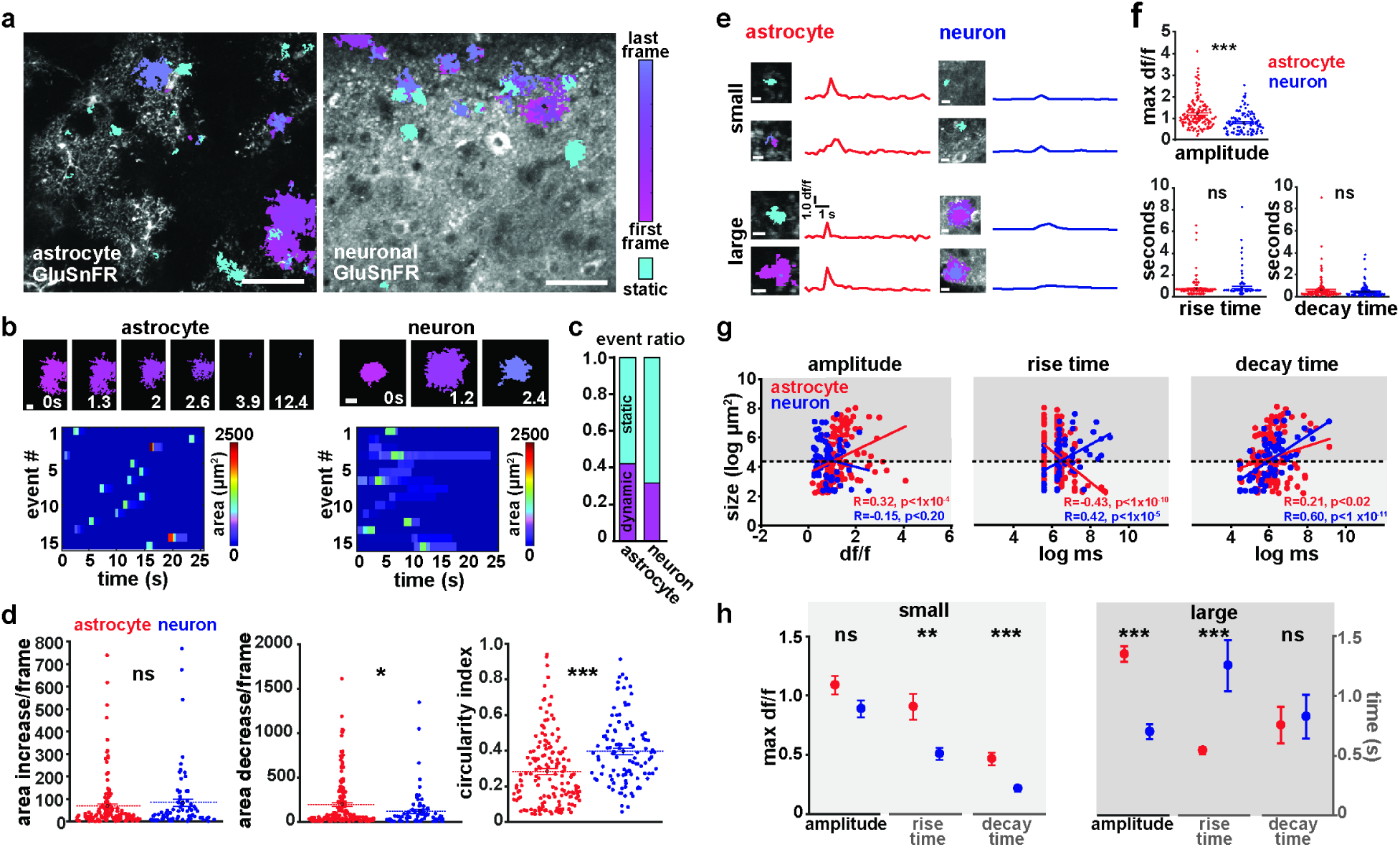
Extracellular glutamate event detection reveals differences between astrocytic and neuronal expression of GluSnFR. (**a**) Representative images of *ex vivo* slices with expression of astrocytic (left) or neuronal (right) GluSnFR. Color indicates detected events. Those with dynamic shape are shown in magenta, and static events in cyan. Scale bar = 50μm. **(b)** Examples of timecourse of astrocytic (left, top) and neuronal (right, top) glutamate events. Scale bar = 10μm. Raster plot of area of astrocytic (left, bottom) and neuronal (right, bottom) glutamate events (n = 15/cell type). **(c)** Ratio of events that change shape over time (magenta) to events that do not (cyan). **(d)** Size dynamics (area increase [left] and decrease [middle] per frame) and shape (circularity index, right) of glutamate events when GluSnFR is expressed on astrocytes (red) or neurons (blue). Same color scheme continues throughout figure. **(e)** Representative small and large glutamate events and corresponding dF/F traces. Scale bar = 10μm. **(f)** Amplitude (top), rise time (bottom left), and decay time (bottom right) differences between all astrocytic and neuronal GluSnFR events. **(g)** Correlations between event size and amplitude (left), rise time (middle), and decay time (right). Dashed line represents size threshold dividing small events from large (see Supp. Fig 11f or details). **(h)** Summary quantification of differences in amplitude and dynamics between small (left) and large (right) astrocytic (red) and neuronal (blue) GluSnFR events. Data are shown as mean ± SEM.

Because AQuA is designed to detect events independent of shape or size, events of heterogeneous size and shape were revealed when analyzing the GluSnFR data (Fig. 5a–b). A large proportion of GluSnFR events changed size over the course of the event, with 42% of total astrocytic and 32% of total neuronal glutamate events exhibiting changes in area (Fig. 5c). On average, astrocytic GluSnFR events were significantly larger (274 ± 39.56 μm^2^) than neuronal events (172 ± 57.06 μm^2^), sometimes encompassing an entire astrocyte (Supp. Fig. 10). Neuronal GluSnFR events were significantly more circular (Figure 6b–d), reflecting morphological differences between cell types. We also found that between cell types, GluSnFR events exhibited different size dynamics (Fig. 5b–d). While there was no difference in the rate of increase in event size between astrocytes and neurons, we did observe that the rate of size decrease of astrocytic events between frames was larger than that of neuronal events (Fig. 5d).

To investigate size differences between cell types more thoroughly, we extracted dF/F from each event by calculating the average fluorescence intensity of the maximum area of each event over its lifetime (Fig. 5e). When we compared these curves, we found that the amplitude of astrocytic events was significantly larger than that of neuronal events (1.2 ± 0.05 vs. 0.79 ± 0.05, Fig. 5e, f [top]), whereas the dF/F rise times and decay times showed no significant difference between cell types (Fig. 5f, bottom). However, we noticed a large spread in the amplitude and kinetics of both cell types (Fig. 5f) and next asked whether the size of each event correlates with these features, and whether these correlations could differentially describe each cell type. We observed some significant correlations of event size with amplitude, rise time, and decay time (Fig. 5g). Specifically, astrocytic GluSnFR event size positively correlated with amplitude and decay time and negatively correlated with rise time (Fig. 5g, red). On the other hand, neuronal GluSnFR event sizes were positively correlated with rise and decay times, but showed no significant relationship with amplitude (Fig. 5g, blue). These correlations indicate that variations in dF/F features within each cell type were size-dependent.

To further explore the entire population of glutamate events for each cell type, we set a size threshold to separate small and large events (Figure 5g, black dashed lines, Supp. Fig. 10). When we do this, we find that the larger amplitude observed in astrocytic glutamate events (Figure 5f, *top*) is dominated by significantly higher amplitudes in large-size astrocytic events, whereas small-size events were not significantly different in amplitude between the two cell types (Fig. 5h). In contrast, while we find no significant difference between cell types in rise and decay time for all events (Fig. 5f, *bottom*), we do observe some significant differences in rise and decay times in the small- or large-size groups separately. In fact, when separating out small and large events, we observe opposite rise-time patterns: small astrocytic events have longer rise times than neuronal events, but large astrocytic events have shorter rise times than neuronal events. Together, these data demonstrate that application of AQuA to GluSnFR images uncovered a class of small extracellular glutamate events. Separation of these small events from their larger counterparts revealed temporal differences between GluSnFR expressed on astrocytes and neurons, suggesting that these differences may reflect different cellular, or cell-type, mechanisms that lead to these extracellular glutamate flux.

## Discussion

With the development and application of a powerful, event-based analysis tool for astrocyte imaging datasets, we have opened the door for quantifying observed fluorescence dynamics, including those that are un-fixed, propagative, and vary in size. We demonstrate that AQuA performs better than other image analysis methods—including those designed for astrocytic and neuronal applications—on these types of simulated datasets, and describe previously unknown phenomena in three types of commonly acquired datasets using the genetically encoded GCaMP or GluSnFR indicators. Because AQuA is data-driven, it can be applied to datasets that have not been directly tested here, including those captured under different imaging magnifications and spatial resolutions. In addition, since the AQuA algorithm functions independently from frame rate, datasets captured with faster frame rates^12,26^ are also just as amenable to an event-based analysis with AQuA as those shown here. Further, AQuA is applicable to fluorescent indicators, particularly those that exhibit complex dynamics, other the ones tested here.

We envision the AQuA software and its underlying algorithm as enabling problem-solving for a wide range of astrocyte physiological questions, both because AQuA more accurately captures dynamics exhibited by commonly used fluorescent indicators than other methods and because there are many more features extracted by AQuA that can be analyzed than those extracted by existing methods. In the current work, we use these multiple features to describe the baseline, or spontaneous, astrocyte physiology in a particular neurobiological circuit, but with varying spatial scale, molecular probe, and experimental preparation. In future work, we and others can apply AQuA-based analyses to circuits in other brain regions and layers to describe potential functional heterogeneities across astrocytes^27^. Beyond baseline differences, we expect that AQuA will be a powerful tool to quantify physiological effects of pharmacological, genetic, and optogenetic manipulations, among others. These manipulations and subsequent analyses would allow researchers to examine both astrocyte-intrinsic and -extrinsic physiology, depending on whether astrocytes, neurons, or another brain cell type is being changed.

There remains significant disagreement in the field about basic physiological functions of astrocytes. Perhaps the most outstanding issue is whether astrocytes undergo vesicular release of transmitters such as glutamate. While we don’t address this controversial topic in the current work, we expect that the heterogeneous activities that we uncover using an AQuA-based analysis of GluSnFR may be key in determining different sources of glutamate in neural circuits under different conditions, and could help untangle some of the conflicting data in this arena. Our tool enabled us to identify extracellular glutamate changes not only by cell type, but also by event size and shape dynamics, demonstrating the most in-depth analysis of GluSnFR data—whether astrocytic or neuronally expressed—than ever before. The event-based analytical tools presented here may be particularly useful as next-generation GluSnFR variants become available and make multiplexed imaging experiments increasingly accessible^28^.

As demonstrated by its utility with both Ca^2+^ and glutamate data, AQuA also has the potential to be applied to other fluorescence imaging datasets that exhibit non-static or propagative activity. Although we designed AQuA specifically to study dynamic astrocyte fluorescence, it is user-tunable, and we anticipate that experimentalists will find it advantageous in other contexts in which neuronal or non-neuronal cells exhibit non-static or propagative fluorescence activity. For example, recently described Ca^2+^ activity in oligodendrocytes displays some similar properties to that in astrocytes^29,30^ and AQuA-based analysis may be useful. Likewise, subcellular compartments in neurons, such as dendrites or dendritic spines, have also been shown to exhibit propagative, wave-like signals^31^ and large-scale, whole-brain neuronal imaging can capture burst-like, population-wide events^32^ as observed in astrocytes *in vivo*. While we predict that the potential applications are wide, it is also important to note the limitations of AQuA, and be clear about when it will not be the most effective approach. Since AQuA detects local fluorescence changes as events, it is not well suited to strictly morphological dynamics, such as those observed in microglia, and it does not improve on the many excellent tools built for analyzing somatic neuronal Ca^2+^ activity^14,15^, where ROI assumptions are well satisfied. In addition, AQuA is designed to analyze 2D datasets only, as these comprise the majority of ongoing dynamic imaging experiments. In the future, AQuA can be adapted to accommodate 3D imaging experiments, including those currently being performed in astrocytes^26^.

When surveying dynamic astrocyte imaging data, particularly Ca^2+^ imaging data, experimental regimes can largely be grouped into two categories: single-cell, usually *ex vivo* imaging and population-wide, *in vivo* imaging focusing on large-scale activity of many cells. Experimental data and neurobiological conclusions from these two groups can differ quite widely, or even conflict with each other. This may be due, in part, to the large, population-wide bursts observed with the onset of locomotion *in vivo*. Many techniques used to analyze these bursting events—all ROI-based—can under-sample events that occur between bursts by swamping out smaller or shorter signals. Here, we present a technique that can be used to sample small- and large-scale activity in the same dataset, or across datasets, allowing researchers to bridge spatiotemporal scales robustly in these types of data for the first time. We believe that this event-based analysis tool will enable astrocyte biologists not only to resolve outstanding physiological problems, but also identify and tackle new ones.

## Supporting information

Movie related to the ex-vivo study

Movie related to the in-vivo study

Movie related to the glutamate study

## Acknowledgements

The authors acknowledge members of the Poskanzer and Yu labs for helpful discussions and comments on the manuscript. We thank Gregory Chin for excellent technical assistance. K.E.P. is supported by NIH R01NS099254, NSF 1604544, Brain Research Foundation Frank/Fay Seed Grant, E. Matilda Ziegler Foundation for the Blind, and the Bold & Basic grant from the QBI at UCSF. G.Y. is supported by NIH R01MH110504 and NSF 1750931.

## Author Contributions

K.E.P and G.Y. conceived and designed the study. Y.W. and G.Y. designed and implemented the AQuA algorithm, software, and simulations. T.V. carried out GluSnFR imaging experiments, M.R. performed *in vivo* Ca^2+^ imaging experiments, and M.C. performed *ex vivo* Ca^2+^ imaging experiments. N.V.D. analyzed all imaging data and provided critical conceptual input to software design. X.M. implemented the Java version of the software. K.E.P supervised the experimental team. G.Y. supervised the computational team. Y.W., N.V.D., K.E.P, and G.Y. wrote the manuscript with input from all authors.

## Methods

### Viral injections and surgical procedures

For slice experiments, neonatal mice (Swiss Webster, P0–P4) were anesthetized by crushed ice anesthesia for 3 minutes and injected with 90nL total virus of *AAV5-GFaABC1D. Lck-GCaMP6f, AAV5-GFaABC1D.cyto-GCaMP6f, AAV1-GFAP-iGluSnFR*, or *AAV1-hsyn-iGluSnFR* at a rate of 2–3nL/sec. Six injections 0.5μm apart in a 2×3 grid pattern with 15nL/injection into assumed V1 were performed 0.2μm below pial surface using a UMP-3 microsyringe pump (World Precision Instruments). Mice were used for slice imaging experiments at P10–P23.

For *in vivo* experiments, adult mice (C57Bl/6, P50–P100) were given dexamethasone (5mg/kg) subcutaneously prior to surgery and then anesthetized under isoflurane. A titanium headplate was attached to the skull using C&B Metabond (Parkell) and a 3mm diameter craniotomy was cut over the right hemisphere ensuring access to visual cortex. Two 300nL injections (600nL total virus) of *AAV5-GFaABC1D.cyto-GCaMP6f* were made into visual cortex (0.5–1.0mm anterior and 1.75–2.5mm lateral of bregma) at a depth of 0.2–0.3mm and 0.5mm from the pial surface, respectively. Virus was injected at a rate of 2nL/s, with a 10min wait following each injection to allow for diffusion. Following viral injection, a glass cranial window was implanted to allow for chronic imaging and secured using C&B metabond^33^. Mice were given at least ten days to recover, followed by habituation for three days to head fixation on a circular treadmill, prior to imaging.

### Two-photon imaging

All 2P imaging experiments were carried out on a microscope (Bruker Ultima IV) equipped with a Ti:Sa laser (MaiTai, SpectraPhysics). The laser beam was intensity-modulated using a Pockels cell (Conoptics) and scanned with galvonometers (or resonant scanners). Images were acquired with a 16x, 0.8 N.A. (Nikon, *in vivo*) or 40x, 0.8. N.A. objective (Nikon, *ex* vivo) via a photomultiplier tube (Hamamatsu) using PrairieView (Bruker) software. For imaging, 950nm (GCaMP) or 910nm (GluSnFR) excitation and 510/84 emission filter was used.

### Ex vivo GCaMP and GluSnFR imaging

Coronal, acute neocortical slices (400μm thick) from P10–P23 mice were cut with a vibratome (VT 1200, Leica) in ice-cold cutting solution (in mM): 27 NaHCO_3_, 1.5 NaH_2_PO_4_, 222 sucrose, 2.6 KCl, 2 MgSO_4_, 2 CaCl_2_. Slices were incubated in standard continuously aerated (95% O_2_/5% CO_2_) artificial cerebrospinal fluid (ACSF) containing (in mM): 123 NaCl, 26 NaHCO_3_, 1 NaH_2_PO_4_, 10 dextrose, 3 KCl, 2 CaCl_2_, 2 MgSO_4_, heated to 37°C and removed from water bath immediately before introducing slices. Slices were held in ACSF at room temperature until imaging. Experiments were performed in continuously aerated, standard ACSF. 2P scanning and acquisition were carried out at 1.06Hz at 512 × 512 pixel resolution. For TTX experiments, 0.5μM of Tetrodotoxin Citrate (Hello Bio) was added to aerated, standard ACSF. 8 minutes elapsed before resuming imaging.

### In vivo GCaMP imaging

At least two weeks following surgery mice were head-fixed to a circular treadmill and astrocyte calcium activity was visualized at ~2hz effective frame rate from layers 2/3 of visual cortex with a 512×512 pixel resolution at 0.8 microns/pixel. Locomotion speed was monitored using an optoswitch (Newark Element 14) connected to an Arduino.

### AQuA algorithm and event detection

#### Overview of the AQuA algorithm

Astrocytic events are heterogeneous and varying with respect to many aspects of their properties. In AQuA, we extensively applied machine learning techniques to flexibly model these events, so that our approach is data-driven and physiologically relevant parameters are extracted from the data instead of imposing *a priori* assumptions. Probability theory and numerical optimization techniques were applied to optimally extract fluorescent signals from background fluctuations. Here, we first delineate the eight major steps in AQuA (Supp. Fig. 1) and then describe key technical considerations in further detail.

Step 1: data normalization and preprocessing. This step removes experimental artifacts such as motion effects, and processes the data so that noise can be well approximated by a standard Gaussian distribution. Particular attention is paid to the variance stabilization, estimate of baseline fluorescence, and variance. Step 2: detect active voxels. Step 3: identify seeds for peak detection. Step 4: detect peaks and their spatiotemporal extension. These three steps work together to achieve peak detection. To detect peaks we start from a seed, which is modeled as a spatiotemporal local maximum. However, since random fluctuations due to background noise can also result in local maxima, we need to detect active voxels such that only the local maxima on the active voxels are considered as seeds. Here, active voxels are those likely to have signals. Step 5: cluster peaks to identify candidates for super-events. Temporarily ignoring the single-source requirement, the set of spatially-adjacent and temporally-close peaks is defined as a super-event. However, clustering results of spatially adjacent peaks are not super-events themselves, because a peak group may consist of noise voxels and temporally distant events. Step 6: estimate the signal propagation patterns. Step 7: Detect super-events. To get super-events from peak clusters, we compute the temporal closeness between spatially adjacent peaks by estimating signal propagation patterns. The propagation pattern for each event is also important for its own sake, by providing a new way to quantify activity patterns. Step 8: split super-event into individual events with different sources. A super-event is split into individual events by further exploiting propagation patterns. Based on propagation patterns within a super-event, the locations of event initiation are identified as local minima of the onset time map. Each initiation location serves as the event seed. Individual events are obtained by assigning each pixel to an event based on spatial connectivity and temporal similarity.

##### Step 1(Data normalization and preprocessing)

We correct for motion artifacts in the *in vivo* dataset using standard image registration techniques^21^ before applying AQuA. However, AQuA does not necessarily require motion correction because it performs event-based analysis, which is localized temporally and thus less prone to motion artifacts.

We perform data normalization and preprocessing to approximate noise by a Gaussian distribution with mean=0 and standard deviation=1. To achieve this, we first apply a square-root transformation to the data to ensure that the noise variance after transformation is not related to the intensity itself, an operation also known as variance stabilization. Second, the noise variance of the transformed data for each pixel is estimated as half of the median of the square of differences between two adjacent values in the time series at the pixel. Mathematically, denote *X_i_* the time series at the *i*th pixel, where *X_i_*[*t*] is the value of the *t*th time point. Then, the noise variance 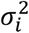 at the *i*th pixel is estimated as

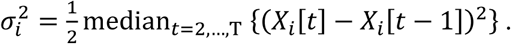

We do not use the conventional sample variance 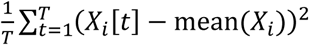 as the estimate. Otherwise, it is inclined to inflate the variance when signals exist in the time series. Third, to estimate the baseline fluorescence *F*_0_ for each pixel, we compute the minimum of the moving average of 25 time-points in a user-specified local time window (default=200 in our experiments). We do not use the full time series to identify the minimum, in order to be robust to image degradation or other long-term trends. Considering the minimum is a biased estimate of the baseline fluorescence, we add a pre-determined quantity to the minimum to serve as the estimate of *F*_0_. Here, the pre-determined quantity depends on the extent of the moving average and the size of the time window, and is found through simulation. Denote *V_i_*[*t*] as the value of *i*th pixel at the *t*th frame in the raw video data. In the following, all analysis is performed on the normalized data,

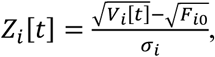

where the subscript *i* denotes that the baseline fluorescence and noise variance are location- and pixel-specific.

##### Step 2 (Detecting active voxels)

A voxel is defined as a pixel of a certain frame. For example, voxel (*x, y, t*) denotes the pixel at location (*x, y*) in the *t*th frame in the movie. An active voxel is the voxel that contains an activity signal. If a voxel is not associated with any event, it is not considered active. Since an event often occupies multiple pixels and extends several frames, we first apply 3D Gaussian filtering to smooth the data to reduce the impact of noise. Then, we calculate the z-scores for each voxel in the smoothed data. Here, z-score is computed as the value of the voxel divided by its standard deviation, which can be estimated as in the normalization procedure above, but now on the smoothed data. All voxels that have z-scores larger than a given threshold are considered tentative active voxels. A liberal threshold is used here to retain most signals, often at a z-score of 3. We next calculate the size of groups of connected tentative active voxels, with spatially connected tentative active belonging to the same group and a minimum size threshold (often 4). If a group of tentative active voxels is less than the threshold, all voxels in this group are removed, resulting in a final list of active voxels.

##### Step 3 (Identifying seeds for peak detection)

Similar to the detection of active voxels, we apply 3D Gaussian smoothing to the normalized data and then find all local maxima, defined as connected components of pixels with a constant intensity value and with all neighboring pixels having a lower value. Considering our time-lapse images as three-dimensional arrays (2D space plus 1D time), each single pixel has 26 neighbors. Although each local maximum generally occupies one pixel due to random fluctuation inherent in the data, this definition allows a local maximum to occupy multiple pixels of the same intensity value. This is helpful for the rare case in which some pixels have saturated values. Because pure random fluctuation can also lead to local maxima, we restrict the search of local maxima to active voxels only. The resultant local maxima are considered seeds for the purpose of peak detection, the subsequent step in the algorithm.

##### Step 4 (Detecting peaks and their spatiotemporal extent)

We partially and temporally extend each seed detected above to all voxels that are potentially associated with each event. We call the seed and its potential extended voxels the *super voxel*. Seeds are processed one-by-one, with higher intensity seeds processed first. Each seed is first extended temporally, then spatially.

The spatiotemporal index (*x*_0_, *y*_0_, t_0_) denotes the seed. When we temporally extend the seed backwards and forwards (Supp. Fig 2b), we encounter two main scenarios. In the first, a voxel before (*x*_0_, *y*_0_, *t*_0_) has a value close to the baseline *F*_0_, and a voxel after (*x*_0_, *y*_0_, *t*_0_) also is close to *F*_0_. If a voxel has an intensity <20% of the seed value, it is defined as close to baseline. In this scenario, the seed is extended temporally until it reaches these two voxels. In the second scenario, extension in either direction never meets a voxel with value that is considered close to the baseline before meeting another seed. To determine whether we these two seeds should be merged, we denote *V*_min_ the minimum value between the two seeds and calculate the difference between *V*_min_ and value at the seed (*x*_0_, *y*_0_, *t*_0_). If the difference is larger than the threshold Δ*_tw_*, which is 2σ_0_ by default for most data, the minimum is considered the end of the extension. Otherwise, these two seeds are merged and the extension continues. For very high peaks, this threshold is too low for perceptually meaningful separation. To split two adjacent high Δ*F* peaks, a strong decrease between them is needed and the threshold is changed to Δ*_tw_* = max(0.3*ΔF*(*x*, *y, t_0_*), 2σ_0_).

Once each seed *i* is temporally extended from (*x_i_*, *y_i_*, *t_i_*) to a peak (*x_i_*, *y_i_*, (*t_i_* − *a_i_*): (*t_i_* + *b*=*_i_*)), (*t_i_ − a_i_*): (*t_i_* + *b_i_*) denotes a time window spanning from *t_i_ − a_i_* to *t_i_ + a_i_*. We define reference curve *c_i_* as the average of nine pixels around (*x_i_*, *y_i_*) in that time window. To spatially extend each peak to cover most signals-of-interest, each seed becomes a set of voxels 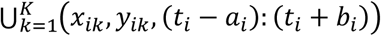 after extension. The corresponding spatial footprint, *K* pixels {(*x_ik_*, *y_ik_*), *k* = 1 … *K*}, is spatially connected. During this process, each seed is associated with two sets. The first is Ω*_i_*, which are pixels already associated with seed *i*. Each pixel in this set, (*x_i_*_1_*, y_i_*_1_), e. g., corresponds to a set of voxels: (*x_i_*_1_*, y_i_*_1_*, t_i_* − *a_i_*:*t_i_ +b_i_*). The second set is Θ*_i_*, the set of pixels to avoid. Initially, Ω*_i_* = {(*x_i_, y_i_*)} and Θ*_i_* is empty.

The spatial extension operation for each seed is repeated a maximum of 40 rounds. For each seed, Ω*_i_* is spatially dilated with a 3×3 square, thus only testing pixels adjacent to the Ω*_i_* boundary. Next, Θ*_i_* is removed from the dilated region. We then test whether each new pixel should be added to Ω*_i_* or not. Because for each given new pixel and each time window we have a time series, we can calculate the Pearson correlation coefficient between this time series and *c_i_*. The correlation coefficient is converted to a z-score using the Fisher transform. If the z-score is higher than the user-defined given threshold, the pixel is added to Ω*_i_*. Otherwise, it is added to Θ*_i_*. Because all seeds are local maxima, no time alignment is needed here.

During the extension process, different super voxels can meet. We want to stop the extension process of one super voxel only when it meets the bright part of other super voxels (50% rising to 50% decaying). For example, we have two peaks from two seeds: (*x*_1_*, y*_1_, (*t*_1_ *− a*_1_): (*t*_1_ *+ b*_1_)) and (*x*_2_*, y*_2_, (*t*_2_ *− a*_2_): (*t_2_* + *b*_2_)). Assume the first seed has already occupied pixel (*x*_3_*, y*_3_). When the second seed tries to determine whether it should extend to (*x*_3_*, y*_3_) or not, we calculate whether (*t*_1_ *− a*_1_*: t*_1_ *+ b*_1_) and (*t*_2_ *− a*_2_*: t*_2_ *+ b*_2_) sufficiently overlap. Two peaks sufficient overlap if the 50% rising to 50% decay ranges of the two peaks overlap. Thus, if (*t*_1_ − *a*_1_*: t*_1_ *+ b*_1_) and (*t*_2_ *− a*_2_*: t*_2_ *+ b*_2_) sufficiently overlap, seed two will not include pixel (*x*_3_*, y*_3_) and it is added to Θ_2_. Otherwise, it is added to Ω_2_. After spatial extension is complete, we remove super voxels with Ω*_i_* < 4 pixels or total voxels < 8 pixels.

##### Step 5 (Clustering peaks to identify candidates for super-events)

A super-event is defined as a group of events connected spatially but originating from different initiation locations. One example is a large burst in the in vivo dataset, where multiple events start at different places but at similar time. Another example is a set of two events originating from different places, propagating and meeting each other in the middle. Thus, in a spatial direction, we may encounter multiple events within the super-event. However, we never encounter two or more events in the temporal direction. To identify candidates for super events, we next cluster peaks, but these results are identical to super-events, because voxels extended to be associated with the peak may have some errors. As discussed below, the candidate super-event must be purified to resolve the final super-event.

Since each super-voxel extends from its seed (representative peak), we also call the process of clustering super-voxels as clustering peaks for conceptual convenience. If two super-voxels are connected and their rise-time difference is less than a given threshold (as discussed below), they are considered neighbors. For two super-voxels, if 10% of pixels of either super-voxel is also occupied by the other, they are a conflicting pair. For each super-voxel, we list all its neighbors and conflicting counterparts. To cluster peaks/super-voxels (Supp. Fig. 2b), we begin with the earliest occurring super-voxel and check each of its neighbors. If a neighbor is not conflicting with that super-voxel, it is combined with the super-voxel. This process is repeated until no new super-voxel can be added. Then we move to the next earliest super-voxel that is not added to any others, and repeat this process. An iterative approach prioritizes events that are close to each other. Supposing the largest rising time difference for super-voxels that is allowed to be neighbors is 10, we start the procedure with the allowed difference as 0 and merge the super voxels. Then we increment the allowed difference by 1 and repeat the step above, until the rising time difference allowed reaches 10.

##### Step 6 (Estimating signal propagation patterns)

For each spatial location/pixel, an associated time series indicates the signal dynamics. Estimation of propagation patterns is formulated as a mathematical problem of time alignment between the time series at each location and a representative/reference time series. Time alignment results directly relay the delay of a given pixel at a time frame with respect to the representative dynamics. Conventionally, time alignment is accomplished by dynamic time warping (DTW)^34^. However, DTW is notoriously prone to noise, which leads to unreliable propagation estimation. Since two adjacent pixels have more similar propagation patterns than two distant pixels, we impose a smoothness constraint on neighboring pixels using our recently developed mathematical model— Graphical Time Warping (GTW)—to explicitly incorporate this constraint^13^. However, since we do not have a representative time series at the very beginning, our strategy is to guess a reference time series from the data and align time series at each pixel to this reference. Then, we use alignment results to obtain an updated reference, and iterate the process of alignment and update of reference until it converges (Supp. Fig. 2b).

To initialize the reference time series, we search for the voxel with the largest Δ*F/F* value and record that voxel’s location. The initial reference is then estimated as the average time series of the pixels in the 5×5 square around that location, with the square size a user-tunable parameter. The voxel with the largest Δ*F/F* value is used because it has the best signal-to-noise ratio. We do not use the time series at a single pixel to initiate the reference because it is noisy, nor do we use the average time series over all the pixels, because the average would be a large distortion to the representative dynamics due to signal propagation. Next, we supply the neighborhood graph and the reference time series to GTW to calculate the time alignment between all pixels and the reference. For each pixel, we consider the 8 pixels around the 3×3 grid as neighbors. A GTW parameter controls the balance between fitness and smoothness of the alignment. We empirically found 1 to be a good value. To control computational complexity, GTW has another parameter corresponding to the maximum time delay allowed. In all our experiments, we found no time delay induced by propagation is larger than 11. So, we set that parameter to 11.

##### Step 7 (Detecting super-events)

Once the time alignment between the representative dynamics and the time series at each pixel is obtained, we refine candidate super-events to obtain final super-events. A super-event is defined as detected when the representative dynamics and all voxels are associated with the super-event. Since each voxel is jointly specified by spatial location and time frame, we next determine which pixels and time frames jointly belong to the super-event. Since representative dynamics are already obtained in the previous step of propagation estimation, here we focus on how to determine which pixels and which time frames are covered by the super-event.

Because each pixel corresponds to a time series, if a pixel belongs to a super-event, its time series should be highly correlated to the representative dynamics of the super-event. Note that the correlation is calculated based on the aligned time series to account for the time distortion due to signal propagation. Thus, we first obtain a new time series for each pixel based on the time alignment obtained previously. Then, we calculate the Pearson correlation between each new time series and the representative dynamics, leading to a correlation map. We further convert the Pearson correlation to z-score using Fisher’s transform. Here, we do not use a threshold for each z-score to determine whether that pixel is statistically significantly associated with the super-event because that ignores the neighborhood information in the correlation map and is less statistically powerful. Instead, incorporating the information from the neighboring pixels, we apply our recently developed order-statistics-based region-growing method to determine which pixels should be associated with the super-event (Supp. Fig. 2b)^35^.

To determine which time frames are associated with the super-event, we now examine the representative time series, calculating the maximum intensity along the curve and considering all time frames with intensity >10% of the maximum to be associated with the super-event. Different pixels may have different time frames associated with the super-event. We use the time alignment results above to identify the time frames associated with the super-event for each pixel. A time frame at a given pixel is associated with the super-event as long as its corresponding time frame in the representative curve is associated with the super-event.

##### Step 8 (Splitting super-events into individual events with different sources)

For each super-event, we have a 2D map of rise-time for each pixel by re-aligning the super-event using GTW. The local minima in this map are potential originating locations for events in this super-event. However, noise may produce random local minima, which do not correspond to true originating locations and are removed by merging with spatially adjacent local minima. We use rise-time to determine whether two local minima should be merged. This idea can be illustrated with the following 1D example: [1 2 4 2 2]. The two local minima are the first and the last pixel (pixel *i* and *j*, respectively), occurring at time 1 (*t*_0_) and 2 (*t*_1_), respectively. To determine whether they should be merged, we find all paths connecting them. In this example, there is only one path and the pixel with the latest rise-time in this path is the third pixel (rise-time=4 (*t_m_*)). The distance between pixel *i* and pixel *j* induced by this path is therefore defined as max(*t_m_* − *t*_0_*, t_m_ − t*_1_). If the distance induced by any path is less than the given threshold, these two local minima are merged.

We next separate super-events into individual events by simultaneously extending all remaining local minima. Each remaining local minimum corresponds to one event. Pixels attached to a local minimum are defined as growing. With each iteration, we add the earliest-occurring pixel to a growing event. If the pixel under examination is adjacent to a growing event, it is done, and then we find the next earliest occurring pixels. Otherwise, we add it to the waitlist and continue with the next earliest occurring one. Each time a pixel is successfully added to a growing event, pixels in the waitlist are checked as to whether they can be added to growing events. When the growing process ends, all individual events are obtained.

### Generation of simulation data sets

#### Spatial footprint templates

We built a set of templates for event footprints from real *ex vivo* data which serve as the basis for the ROI maps in the subsequent step. Footprints are processed by morphology closing, hole filling, and morphology opening to clean boundaries, with 1683 templates generated total.

#### ROI maps

2D ROI maps generated from spatial footprint are used to generate events in subsequent steps. Different simulation types have a different preference for the size of the ROIs. Maximum number of ROIs is set at 100; ROIs are randomly chosen and placed onto a 2D map <5 pixels from existing ROIs.

#### Simulation dataset 1 (size-varying events)

To simulate event size changes, we generate events for each ROI and then alter them to have different sizes so that each ROI in the 2D map will be related to multiple events whose centers are inside that ROI, but whose sizes are different. The degree of size change is characterized by the odds ratio (maximum = 5) between the maximum and the minimum allowable sizes of the events associated with that ROI. For example, with an odds ratio of 2, the size of the event will range from 50–200% of the ROI area. The chances for the event size to be larger or smaller than the area of the ROI are the same. To achieve this, we generate a random number between 1 and 2, then randomly assign whether to enlarge size by multiplying or shrink by dividing by this factor. Event duration is four frames.

To determine the frames at which the event occurs, we first put the event 10–30 frames (randomly) after the ROI occurs. Spatial distance of this event from others must be ≥3 pixels and temporal distance ≥4 frames. Part of the event may be inside the spatial footprint of other ROIs, as long as its spatiotemporal distance to other events is larger than the threshold set above. Events are generated for each ROI; on average, we simulate 250 frames with 800 events on 90 ROIs.

#### Simulation dataset 2 (location-changing events)

To simulate event location changes, we generate events with the same size for each ROI and shift them to nearby locations. Thus, each ROI (450–550 pixel size) is related to multiple events near to that ROI. Denote *dist* the distance between the event center and the ROI center. Denote *diam* the diameter of the ROI. The degree of location change is quantified by the ratio between *dist* and *diam*. For example, if we set 0.5 as the maximum degree of location change, the distance of the center of a new event to the ROI will be 0–0.5 times the diameter of the ROI. If the ratio is 0, we simulate a pure ROI dataset. The new event may be located any direction from the ROI, randomly picked from 0–2π. Shapes of new events are randomly picked from the templates, so may be different from the ROI while size is constant. Event duration is four frames, and the remaining steps are the same as above. On average, we simulate 250 frames with 800 events on 90 ROIs.

#### Simulation dataset 3 (propagating events)

We simulated two types of propagation: growing and moving, leading to three types of synthetic datasets: growing only, moving only, and mixed. These three types are generated similarly. The ROI map is generated as above, and ROI sizes are 4,000–10,000 pixels, with events generated inside each ROI. In comparison, events in the size-change and location-change simulations can be (fully or partially) outside their corresponding ROIs. We simulate only one seed (starting propagation point) in each ROI. For each event, we generate a rise-time map (for each pixel in the ROI) and construct event-propagation based on the map. We obtain this map by simulating a growing process starting from the seed pixel, with the seed pixel active at the first time-point. At the next time point, its neighboring pixels are active with a variable success probability. Growth continues until ≥90% of pixels in the ROI are included in the event. Based on the rise-time map, we identify frames at which pixels become active in the event. To determine when the event ends, we treat growing and moving propagation differently. In growing propagation, all pixels are inactive simultaneously 2 frames after the last pixel becomes active. For moving propagation, the duration is 5 frames. Typically, we generate approximately 140 events in 14 ROIs for each synthetic dataset.

#### Simulate various SNRs

Gaussian noise is added to the synthetic data to achieve various SNRs. We define the signal intensity as the average of all active pixels in all frames. SNR is defined as

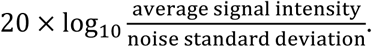

When we change the degree of location change, size change, and propagation duration, we add noise with 10 dB SNR. To study the impact of SNR on size changes, size-change degree is 3. For location changes, distance-change ratio is 0.5 while varying SNRs. For propagation, propagation duration is 5 frames. Seven SNRs are tested: 0, 2.5, 5, 7.5, 10, 15, 20 (all in dB).

#### Post-processing simulated data

We set the average signal intensity at 0.2, with a range from 0–1. Synthetic data is spatially filtered to mimic blurred boundaries in real data. The smoothing is performed with a Gaussian filter with a standard deviation of 1. Signals with intensity <0.05 after smoothing are removed. Remaining signals are temporally filtered with a kernel with a decay τ of 0.6 frames. The rising kernel is linear. For propagation simulation, data is down-sampled by five. Next, we perform a cleaning step. For each pixel in each event, we find the highest intensity (x_peak) across frames. For that pixel, we set signals that are <0.2 times of x_peak to 0. Finally, a uniform background intensity of 0.2 is added (except for GECI-quant, where no background is added; see below).

### Application of AQuA and peer methods on the simulation data sets

Based on our knowledge about simulated datasets, we apply specific considerations for each analytical method in order to set optimal parameters for each. In this way, we aim to assess the methodological limit of each method, rather than suboptimal performance due to inadequate parameter-setting. We expect that the performance of the peer methods on simulation data is an overestimate of their performance on real experimental data, because here we take advantage of the ground-truth knowledge, which is not available for experimental astrocyte data.

#### Event detection using peer methods

AQuA and CasCaDe report detected events, while other methods report detected ROIs. For a consistent comparison, we detect events from those methods that use ROIs. Once ROIs are detected, we calculate the average dF/F curve for each ROI, as follows: The curve is temporally smoothed with a time-window of 20. The minimum value in the smoothed curve is considered baseline. Assume the minimum value occurs at time *t_base_*. The baseline is then subtracted to obtain the dF curve. The noise standard deviation σ is estimated using 40 frames around *t_base_*. We then obtain a z-score curve as dF/σ. A large z-score indicates an event; we use a z-score threshold of *z*_0_. The value *z*_0_ is set according to ground-truth knowledge, so that the smallest-size event in the simulation data is detected by this threshold. Denote *x*_0_ and *s*_0_ the peak intensity and the size for the smallest event in the ground truth. We also denote the ground truth noise level as *σ*_0_. Then, the threshold is calculated as,

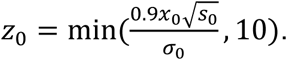

We clip the score to 10 to avoid setting large values for high SNR. For CaSCaDe, we supply this value as the peak intensity threshold parameter.

Using the z-score curves and threshold, we detect events from ROIs for CaImAn, Suite2P, and GECI-quant. For each z-score curve, we find all frames with values *>z*_0_. Each frame is a seed for an event. Assume the z-score for that frame is and we search before and after that frame. If the intensity of the frame is ≥ 0.2*z*_1_, the frame is associated with the event. If we meet frames with intensities < 0.2*z*_1_, we stop searching that direction. Once finished, we obtain all frames associated with the event. We continue with another seed frame to find another event. Note that if a frame is considered part of an event, we do not consider it as a seed for another event, even if it is >*z*_0_. The spatial footprint is fixed for all frames in an event, based on the ROI detected. Combining spatial footprint and frames, we obtain events for each ROI and identify all voxels belonging to an event.

#### Parameter setting for AQuA

The parameters of AQuA are based on the ex-vivo-GCaMP-cyto preset with the following modifications: For different noise levels, we apply different smoothness levels. The smoothing is performed only spatially and values are empirically chosen. The smoothness parameter is the standard deviation of the Gaussian smoothing kernel used.

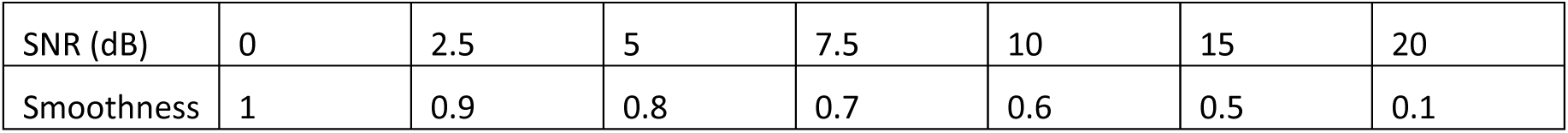

|  |  |  |  |  |  |  |  |
| --- | --- | --- | --- | --- | --- | --- | --- |
| SNR (dB) | 0 | 2.5 | 5 | 7.5 | 10 | 15 | 20 |
| Smoothness | 1 | 0.9 | 0.8 | 0.7 | 0.6 | 0.5 | 0.1 |

We do not simulate motion of the field-of-view, so we do not discard any boundary pixels, and we set regMaskGap = 0. We do not simulate Poisson Gaussian noise; we use additional Gaussian noise only, so PG = 0. Event sizes in the simulation are >200 pixels, so we set the minimum event size to be a value much smaller: minSize = 16. An event may not have more than one peak, so we set cOver = 0. We do not simulate temporally adjacent events, so we set thrTWScl = 4. We do not use proofreading, so we choose a more stringent z-score of events: zThr = 5.

#### Specific considerations for CaSCaDe

We use the following parameters for CaSCaDe: According to the duration and temporal distances of the simulated events, we can safely set peak distance p.min_peak_dist_ed = 3 and minimum peak length p.min_peak_length = 2. We set the spatial smoothing filter size in the 3D smoothing function (bpass3d_v1) according to the size of the event, so we set p.hb equal to 2x median of the radius of the spatial footprint of all events. We use this setting because the default settings could not detect larger events on the simulation data sets. For temporal smoothing, we set p.zhb=21. We do not need to correct background, so we set p.int_correct= 0. The minimum peak intensity is p.peak_int_ed = z0, as discussed above. Minimum event intensity is p.min_int_ed = min(2, p.peak_int_ed *0.2). We modified the low-frequency part of the watershed segmentation step to allow larger events to be detected, by changing the function *bpassW* inside the function *domain_segment*. We replaced the noise estimator in CaSCaDe (function *estibkg*) with the more robust one used by AQuA.

CaSCaDe uses a supervised approach to classify detected events. Instead of manually labeling a large number of events and training many SVM models, we directly use ground truth to perform training. For example, for each event detected by CaSCaDe, we check the ground-truth data to test whether it is (part of) a true event. If so, it is retained; otherwise, it is discarded.

#### Specific considerations for GECI-quant

GECI-quant requires user input at each step. Here, we describe how to automate these steps by taking advantage of ground-truth information. This allows us to test many conditions and repeat many times.

First, we do not add background signals to the synthetic data, so background subtraction is ignored. The domain- and the soma-detection steps require manual thresholding. We estimate the best threshold using the ground-truth data for each simulation. To do so, we scan 255 thresholds and use the one that leads to the best correlation between binarized data and ground truth. We next cleaned the binarized signals with sizes <4 pixels. The data here is also smoothed as it is in GECI-quant (3×3 spatial averaging). Events with spatial footprints < 1,000 pixels are treated as domains and others are treated as somas. The soma segmentation step also uses a threshold. We first process the data as in GECI-quant: for every three frames, a standard-deviation map is calculated so that each voxel in ground-truth data is associated with a standard deviation value. The average of all standard deviations from the ground-truth data is used as the threshold.

We next made the entire analysis pipeline automatic. Fiji is called from the command line in each step and parameters are passed as well. The final ROIs from Fiji are brought back into MATLAB. ROIs are > 15 pixels in area. All other parameters are unchanged, including those for the particle detector. Note that this modification cannot be used as an automated version of GECI-quant for real applications since it relies on ground-truth information.

#### Specific considerations for CaImAn

We experimented with different parameters for CaImAn and found the following set of parameters performed best on simulation data. As event size can be large, we enlarge the patch size, so patch_size = [128,128] and overlap = [32,32]. Components to be found is set to K = 50. The standard deviation of the Gaussian kernel (half size of a neuron) is enlarged to tau = 16. Maximum size is 5,000 and the minimum size is 25. Decay time is 0.5. Other parameters are based on default settings. No spatial or temporal down-sampling is used. Adjusting these parameters dis not impact results on our simulated data. We used the 5/5/2018 version downloaded from https://github.com/flatironinstitute/CaImAn-MATLAB.

#### Specific consideration for Suite2P

The most critical parameter for Suite2P is neuron size. We set db.diameter equal to the minimum between 50 and the median of the radius of the spatial footprint of all events. Setting the diameter too large leads to an out-of-memory issue. We bypass the registration step. We used the 6/4/18 version downloaded from https://github.com/cortex-lab/Suite2P.

### Performance evaluation on the simulated data

To evaluate the accuracy of detected events, we quantify the intersection over the union (IoU). We consider all event voxels, not only pixels as in ROI-based methods. For each detected event *i*, we find all the ground-truth events that have common voxels with event *i*. For each such ground-truth event, e.g., event *j*, we calculate an IoU score (also known as Jaccard index) between this pair of events as the following,

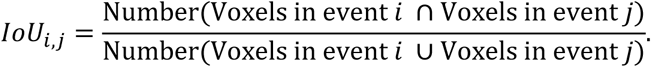

When a detected event can be perfectly matched with a ground-truth event, its IoU score is 1. A score of 0 indicates this pair of events has nothing in common. For each detected event *i*, we find the maximum IoU score among all pairs between this event and a ground-truth event. We denote this maximum score as *IoU_i_*. Similarly, we can compute a score *IoU_j_* for the ground truth event *j*. The final IoU score is obtained by averaging over all events, including detected and ground-truth events. Supposing we have *I* detected events and *J* ground truth events, where *I* and *J* are not necessarily equal, we compute the final score as the following,

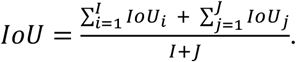

All simulation is performed on a workstation with 16 cores, 128 GB RAM and 6TB hard drive. We use MATLAB 2018a on Windows 10 Enterprise Edition. GECI-quant is run on Fiji with ImageJ version 1.52h. Each simulation is repeated 10 times. The mean and 95% confidence interval (CI) of IoU score is calculated and plotted. The CI is calculated as [*μ_sim_ − 2σ_sim_*, *μ_sim_* + *2σ_sim_*], where *μ_sim_* is the estimated mean and *σ_sim_* is the estimated standard deviation (*σ_sim_*) based on 10 repetitive runs.

### Open-source software for analyzing and visualizing dynamic fluorescent signals in astrocytes

Applying software engineering principles, we developed an open-source toolbox for astrocyte fluorescent imaging data with detailed user guidelines. The software not only implements the AQuA algorithm for detecting events, but also provides an integrated environment for users to see the results, interact with the analysis, and combine other types of data. There are two versions of the software with the same functionality, based on MATLAB or Fiji. The software is freely available at https://github.com/yu-lab-vt/aqua. Detailed documents and example applications can be found there. Here, we highlight several important functions of the software.

First, the software implements AQuA and provides several options to export the event-detection results, including TIFF files with color-coded events, event features in Excel, and MATLAB or Java data structures to be used by other programs. Second, the software can display analysis results by adding color to the raw video, where color encodes the value of a user-defined extracted feature such as propagation speed. Users can specify which feature to be displayed, either an existing feature in AQuA or a user-designed feature based on features provided by AQuA. We provide several pre-defined colormaps, but allow users to manually define colormaps as well. AQuA also provides a side-by-side view, to simultaneously display two features or a raw video plus one feature. Third, the software provides a convenient way to interactively view detected events and their associated features. By clicking on an event, the dF/F curve for the event is shown in a separate panel below the video, and the time-frames during which the event occurs are highlighted in red. The values of several other features for that event are also shown in another panel. The software allows multiple events to be selected simultaneously, so that their curves and features can be plotted together and compared. Fourth, the software provides both automatic and manual ways to proofread the results. For automatic proofreading, events are filtered by setting desired ranges for features-of-interest. Alternatively, users can choose the ‘delete/restore’ button and manually click an event to remove it. Fifth, the software provides flexible ways to incorporate region or landmark information. Users can manually supply regional information such as the cell boundary, or landmark information such as the location of a pipette for pharmacological application. Users can also load region or landmark information from other data sources, such as another fluorescence channel that captures cell morphology. The software can extract landmark-related features for each event, including the direction of propagation relative to a landmark.

**Supplemental Figure 1:**
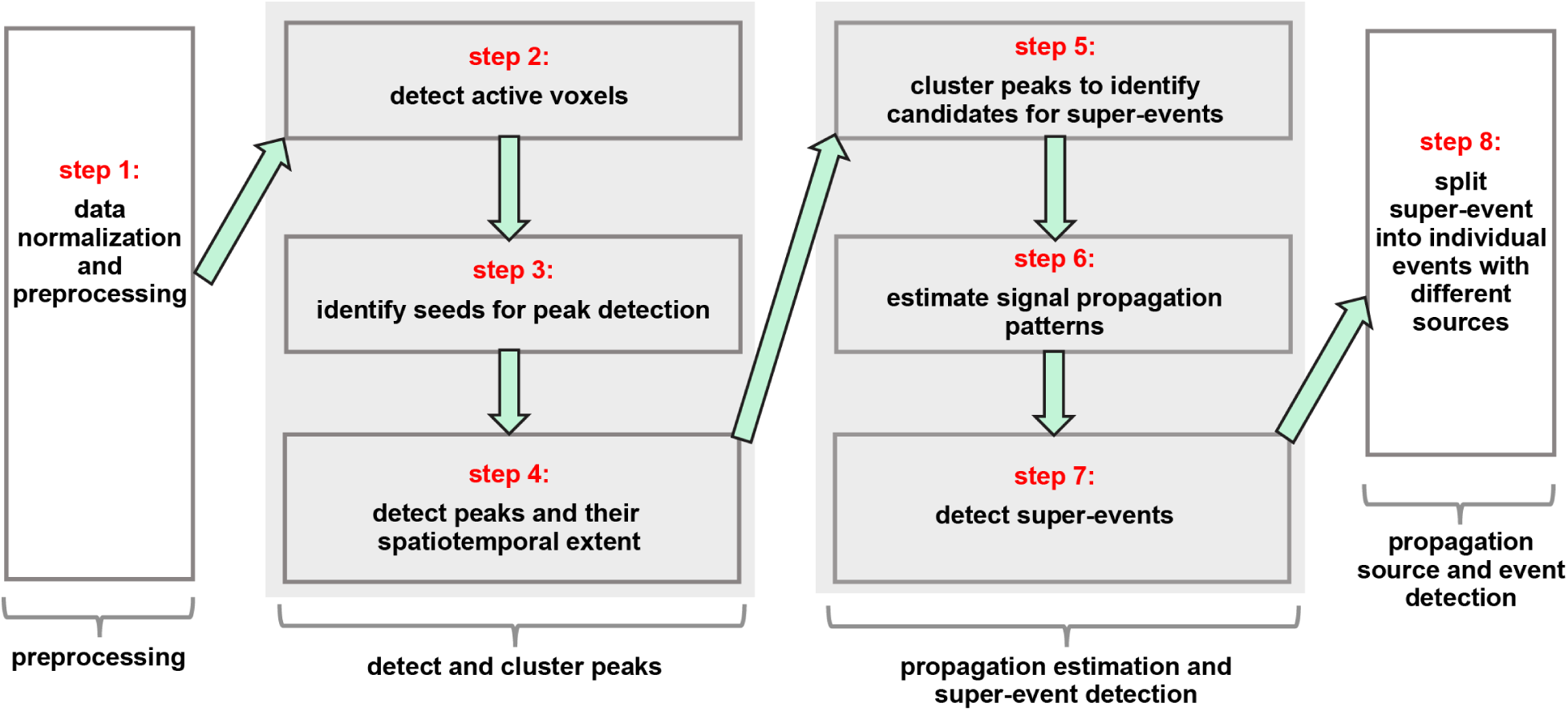
Eight steps in the AQuA algorithm. The eight steps can be grouped into four modules indicated by brackets below panels. The last three modules are further illustrated in Supp. Fig. 2.

**Supplemental Figure 2:**
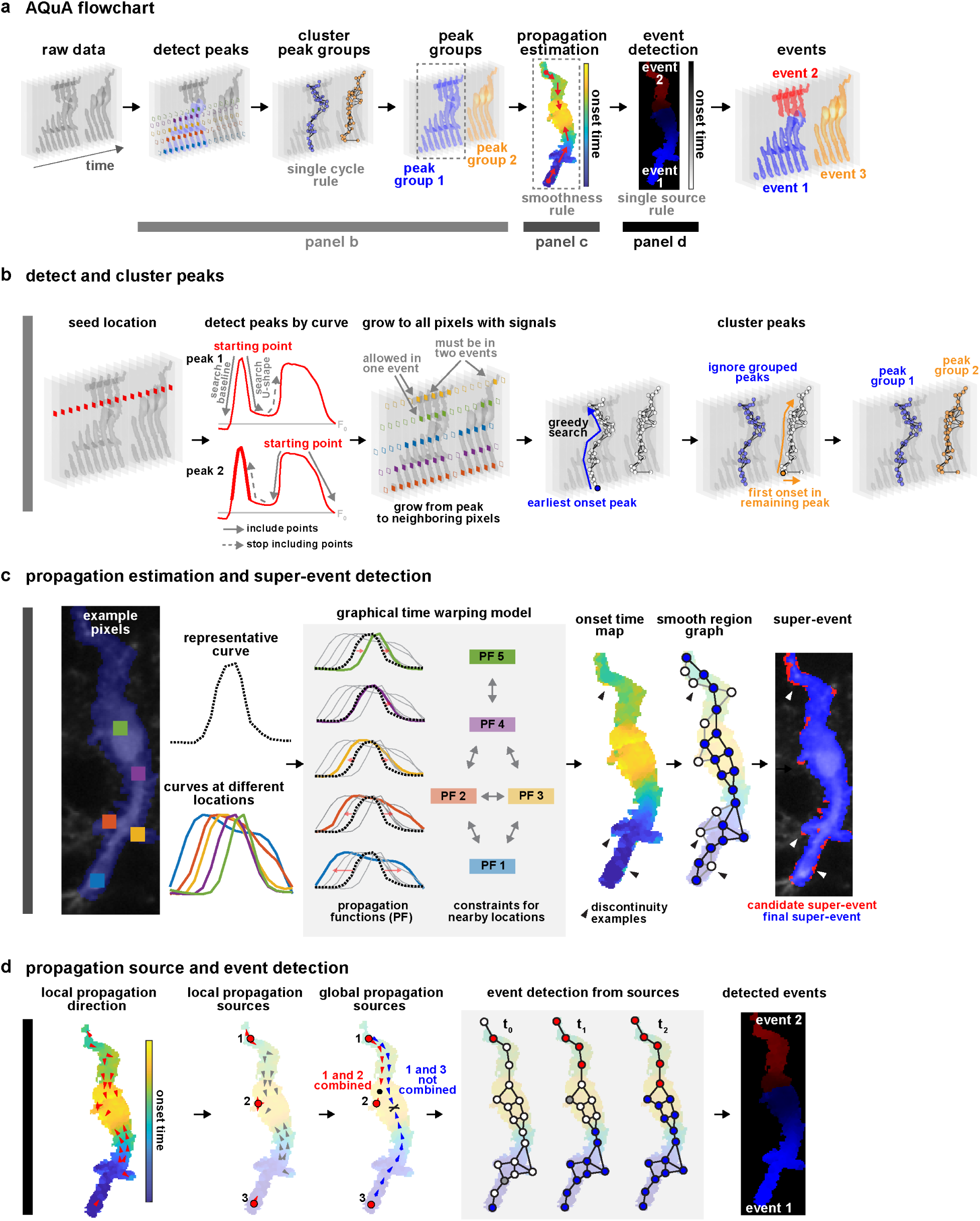
Schematic illustration of three major modules in AQuA algorithm. Curves and regions taken from a real data set. **(a) AQuA flowchart**, with three gray bars below indicating where the three major modules are located with respect to the AQuA flowchart. **(b) detect and cluster peaks**: curves in the *detect peaks by curve* panel are associated with the location labeled by the red diamond in the *seed location* panel. One curve may have multiple peaks, which are detected one-by-one. Once a peak is detected at a seed location, the peak is spatially extended to include its neighboring pixels as in the *grow to all pixels with signals* panel. Clustering of peaks starts from the peak with the earliest onset time and includes its spatially adjacent peaks based on the two inclusion rules shown in the *grow to all pixels with signals* panel. Two peaks at one location are never clustered into one group. Once the greedy search strategy can’t find more peaks to include, it stops and one peak group is formed. Then, to find another peak group, the greedy search restarts from the first onset in the remaining peaks. The process is repeated until no peaks remain. **(c) Propagation estimation and super-event detection:** This module is applied to each peak group. The five colored curves are the dynamics of the five exemplar pixels with corresponding colors. The dashed curve is the representative or reference curve. In the *graphical time warping model* panel, red arrows indicate how the reference curve can be warped to represent the curve at each location. The graphical time warping model incorporates the information that nearby locations should have more similar curves than distant locations. A double-headed arrow between two functions informs the model that these curves should be warped similarly to the reference curve. As a comparison, if there is no double-headed arrow between curves, dissimilar warping functions are allowed. Once the warping function is calculated by the graphical time warping model, onset time is computed for each pixel, resulting in an onset time map. Note discontinuity of onset time examples at black triangles. These pixels are removed to obtain the final super-event, which may contain multiple events and are subject to the next operation. **(d) Propagation source and event detection**: Local propagation sources are obtained by finding local minima on the onset time map. According to the rules described in Methods, some local sources will be combined/merged, resulting in global propagation sources. Briefly, if the path between two local sources does not have to go through a location with a late onset time, these two local sources are combined. Then, each global propagation source leads to an event. Each event is obtained by growing each global source to include its neighboring pixels. In the *event detection from sources* panel, solid dots are pixels already assigned to an event, white dots are unexplored pixels, and grey dots are explored but await a later decision to be assigned to an event.

**Supplemental Figure 3:**
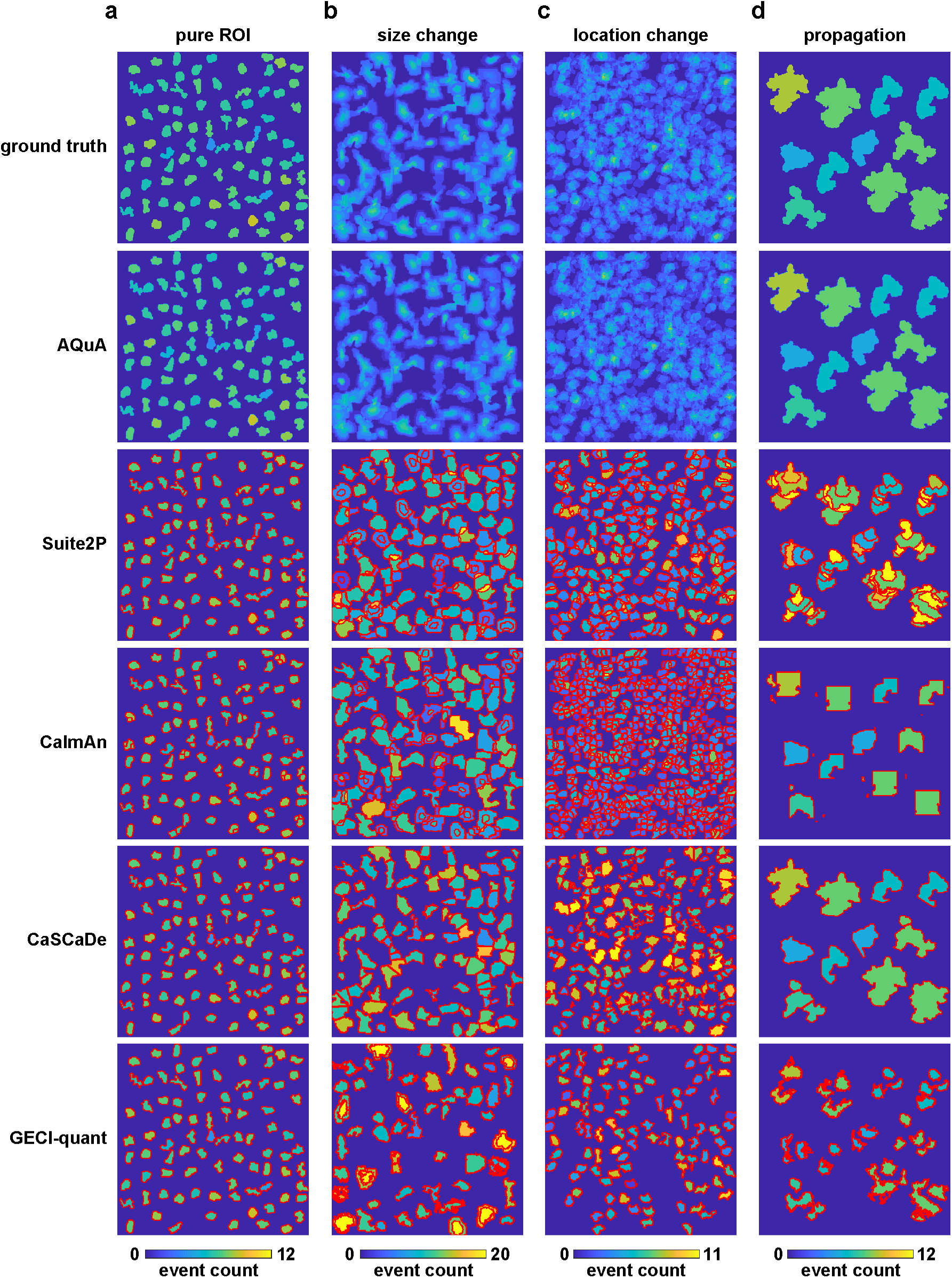
AQuA detects ground truth events across three types of simulated data. Color represents event count for each pixel (note colors bars have different scales in each dataset). Red borders show ROIs detected by ROI-based methods. (AQuA does not detect ROIs.) **(a)** Pure ROI. **(b)** Size change odds of 5, indicating size changes 20–500% of ROI. **(c)** Location change ratio of 1. Average distance to the center of the ROI is 100% the ROI diameter. **(d)** Mixed propagation with 10 frames.

**Supplemental Figure 4:**
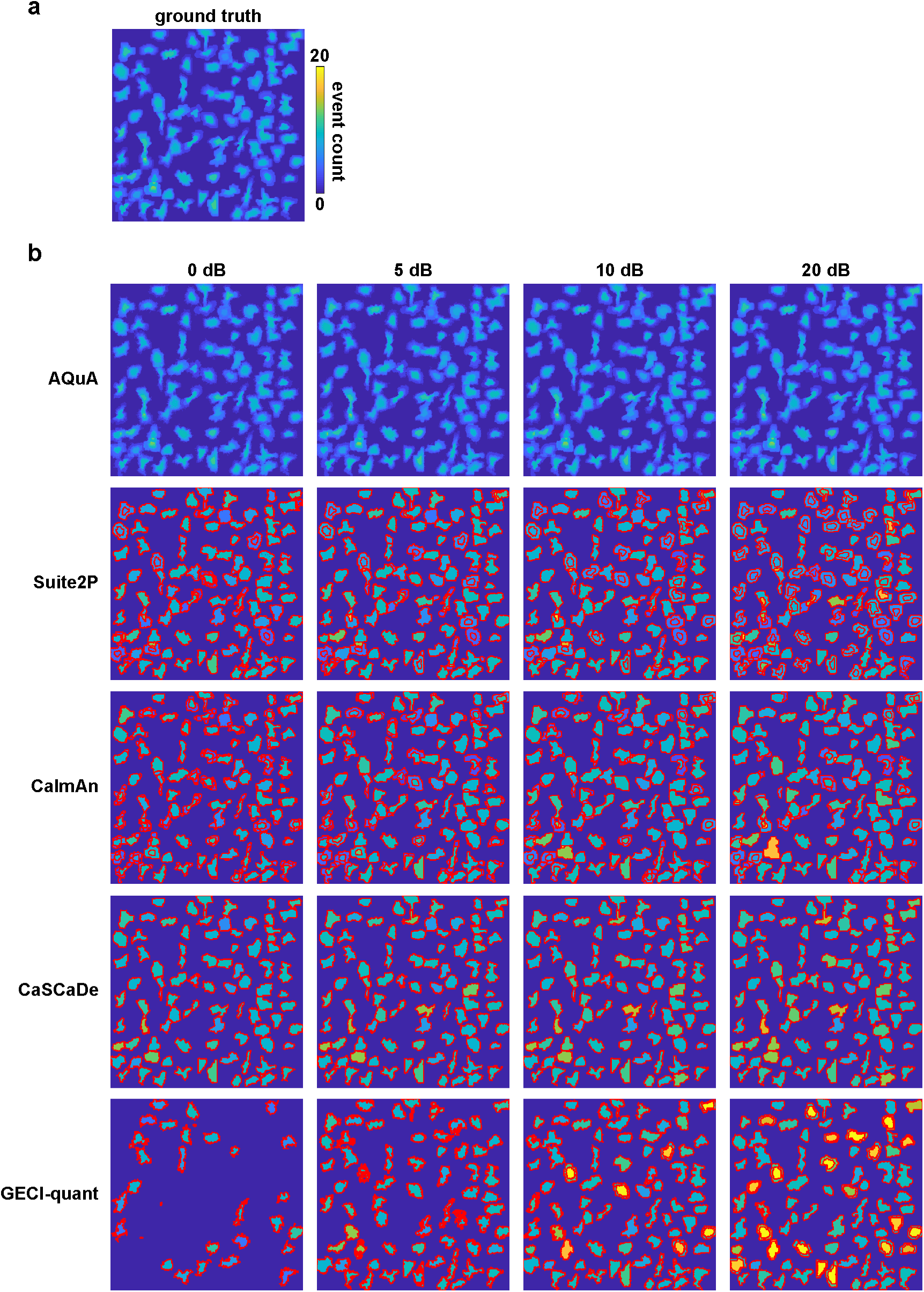
Event counts under different SNRs. Study the impact of SNR change when size change ratio is 3. The color shows the count of events on that pixel. All plots share the same scale. The red lines are the boundaries of detected ROIs. **(a)** Ground truth event count and the color bar for all plots. **(b)** The event count for all methods under four different SNRs.

**Supplemental Figure 5:**
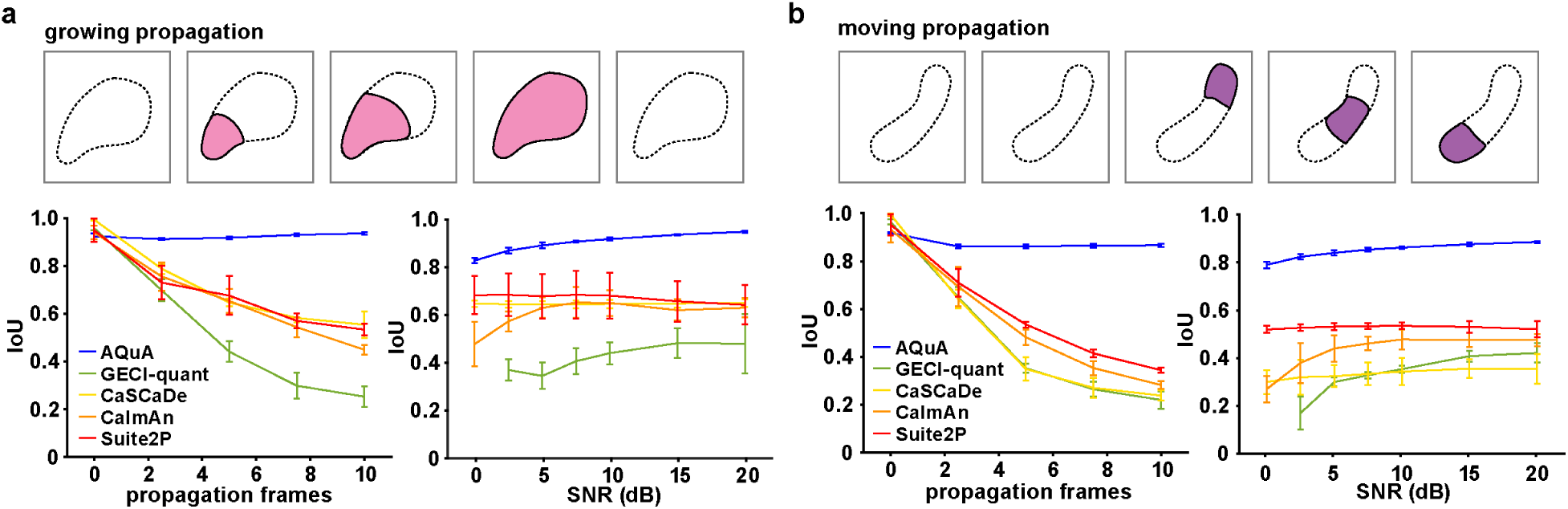
Peer method performance on growing and moving propagation types. Schematic (top) and results (bottom) of performance of five image-analysis methods (AQuA, GECI-quant, CaSCaDe, CalmAn, and Suite2P) on simulated datasets with **(a)** growing propagation and **(b)** moving propagation. Change of the propagation frame number is shown in the bottom left panel, and varying SNR in the bottom right. When the number of propagation frames (not the event duration) is 0, the simulation is under pure ROI assumptions. IoU (intersection over union) measures the overlap between detected and ground-truth events. An IoU of 1 is the best performance achievable by any method, meaning that all detected events are ground-truth and all ground-truth events are detected. The bars on each curve indicate the 95% confidence interval calculated from 10 independent replications of simulation, where each simulation contains hundreds of events.

**Supplemental Figure 6:**
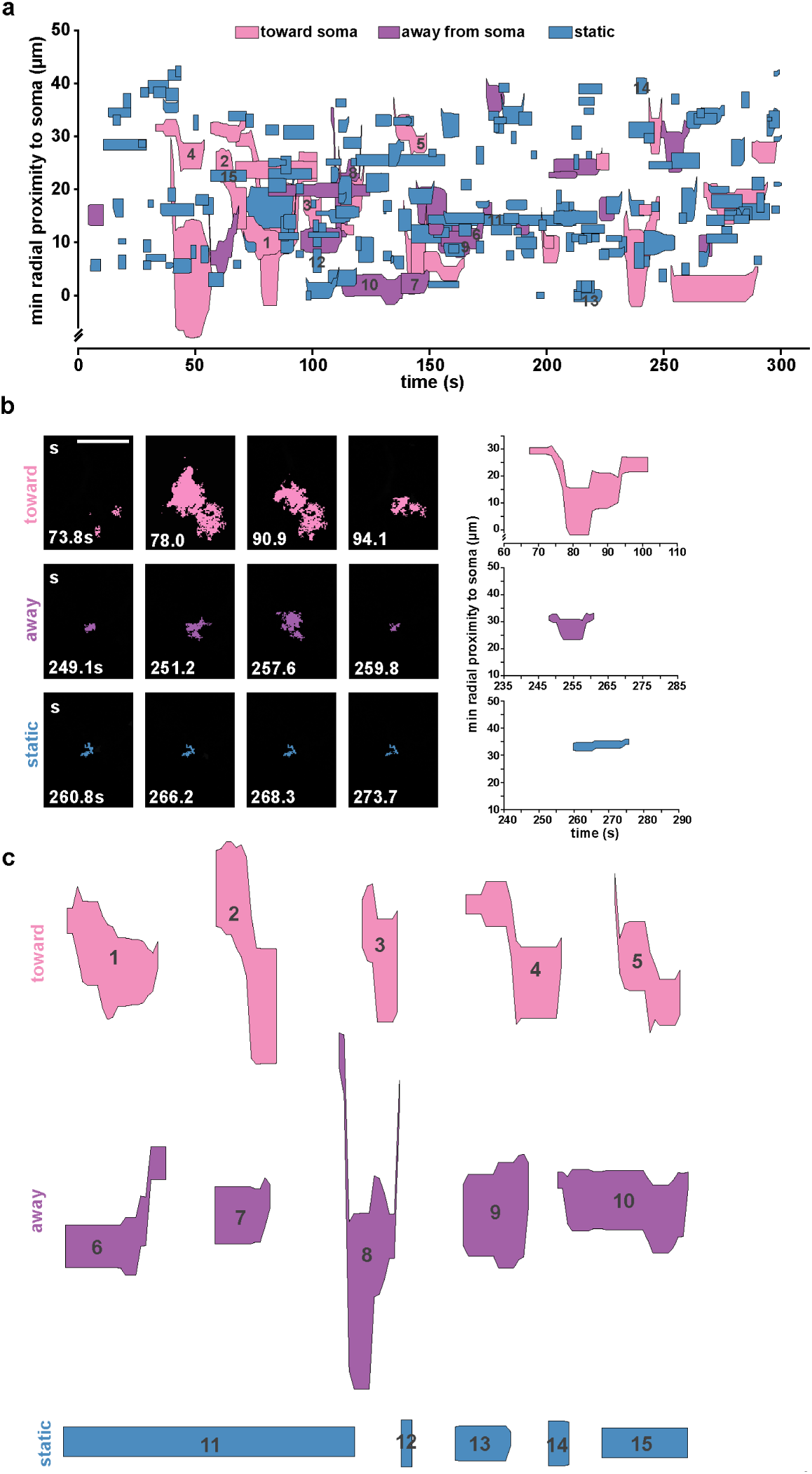
AQuA features enable detailed Ca^2+^ activity plots. **(a)** Spatiotemporal plot of Ca^2+^ activity from a five minute movie (the first minute of which is shown in Fig. 3b). Each event is represented by a polygon that is proportional to the area of the event as it changes over its lifetime, and color-coded by propagation direction. **(b)** Example time series illustrating how propagation direction is determined (left). A propagation direction score is calculated for each event by multiplying the Euclidian distance between the event pixels’ proximity to the soma at each frame by each pixel’s intensity. The overall score is the summation of this weighted pixel intensity distance over the lifetime of the event. Therefore, if more pixels with higher intensity move toward the soma it will be classified as such (top). While some events appear in the plot as moving toward the soma, they are actually calculated as moving away from the soma (middle) since we are only displaying the minimum event proximity to the soma in the spatiotemporal plot, but calculate each pixel’s proximity to the soma when generating propagation score (see Supplemental Movie 4). Further, pixel intensity is first thresholded at 0.3dF/F. Therefore, events that move toward or away from the soma yet have pixel intensities below threshold (bottom) appear to have a propagation direction when plotted, yet have a zero propagation direction score when calculated. **(c)** Additional events plotted for each propagation direction category to demonstrate range of detected/plotted events. Scale is not equivalent to events shown in *b*, but is equivalent within entire group shown here.

**Supplemental Figure 7:**
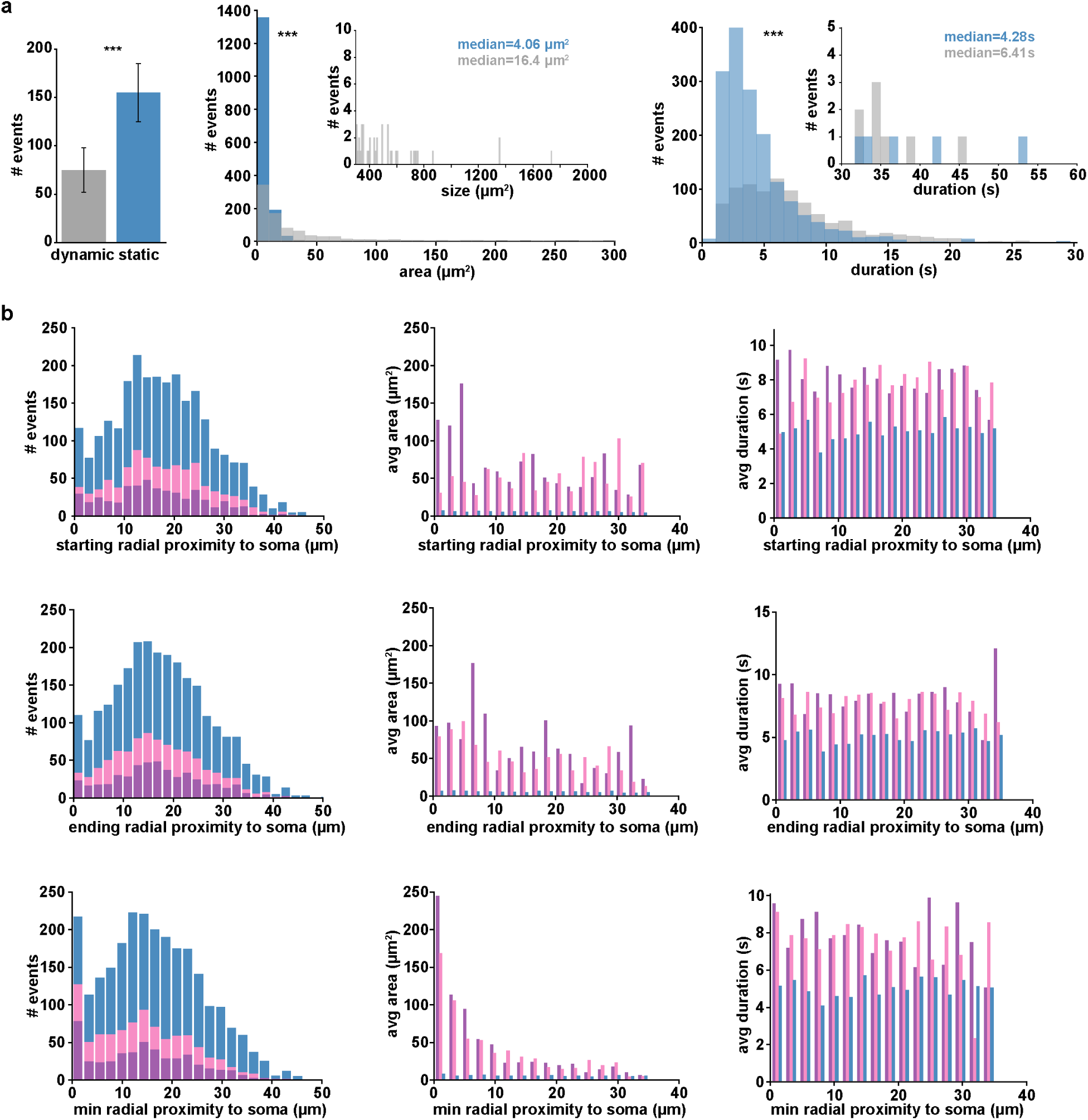
Distribution of Ca^2+^ event features. **(a)** Left: total number of Ca^2+^ events that are dynamic (gray, propagation direction score > 0) and static (blue, propagation direction score = 0), ***p < 0.001, n=11 cells, chi-square test for independence. Middle: distribution of Ca^2+^ event area for dynamic and static events, ***p < 0.001, one-tailed Wilcoxon rank sum test. Right: distribution of Ca^2+^ event duration for dynamic and static events, ***p < 0.001, one-tailed Wilcoxon rank sum test (right). **(b)** Distribution, average area, and average duration of events propagating toward soma (pink), away from soma (purple), and static events (blue) compared to starting distance from soma (top row), ending distance from soma (middle row), and minimum distance from soma (bottom row). Bin widths calculated by Freedman-Diaconis’s rule.

**Supplemental Figure 8:**
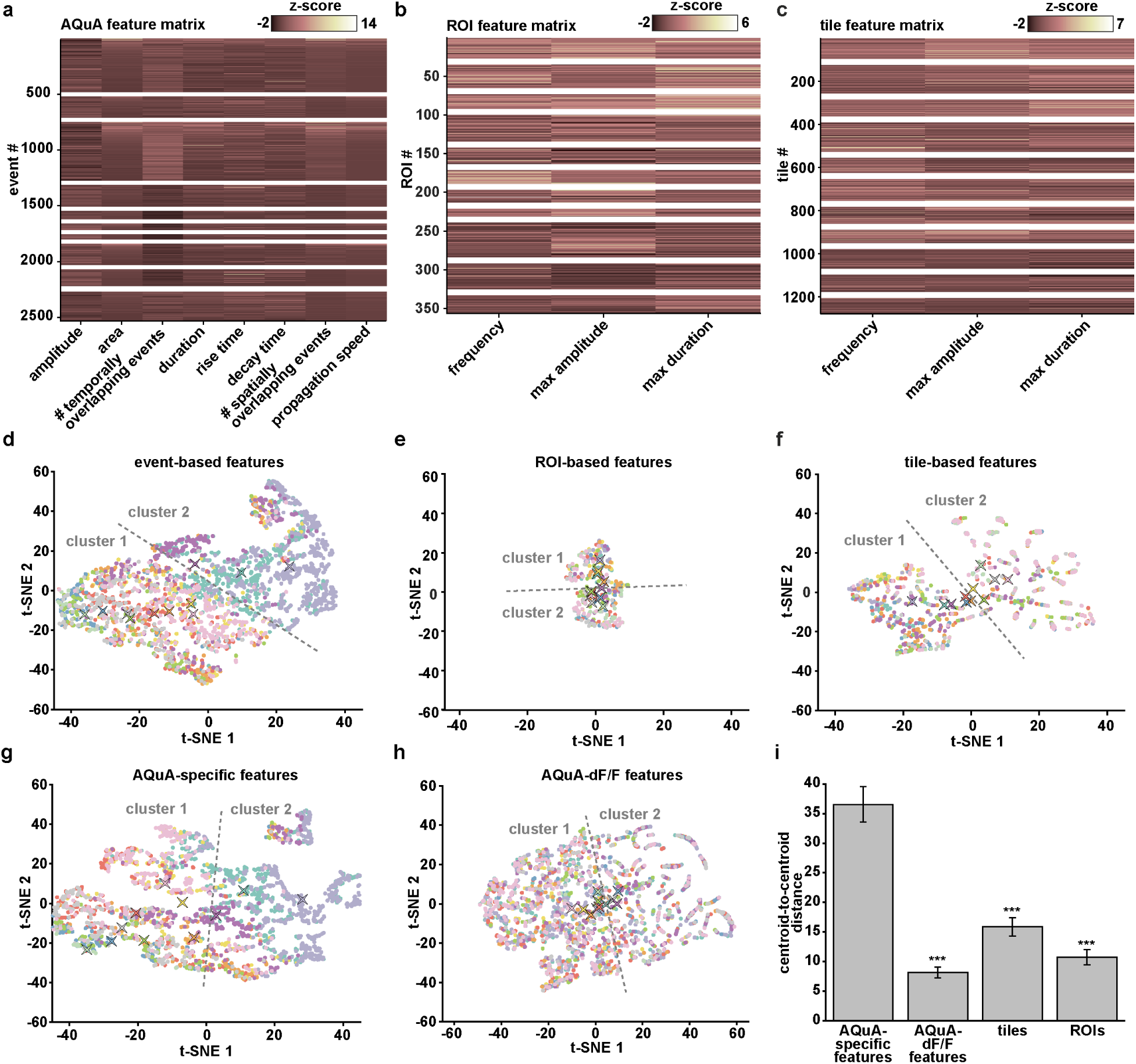
Cluster analysis on features generated from three spatial footprint methods. **(a)** Heatmap of *z*-scores for eight AQuA features (x-axis) describing each event. White boxes demarcate events and features from individual cells. **(b–c)** Top: heatmaps of *z*-scores for three features describing the Ca^2+^ activity at each ROI (*b*) or tile (*c*) location. ROIs detected using average projection image with a 5μm square filter applied (for ROIs) or 5×5μm tiles, based on fluorescence intensity and size. Ca^2+^ events with signals > 0.03dF/F and two times the noise of each individual trace were selected. Pixels within each ROI or tile were averaged and dF/F was calculated by dividing each value by the mean values from the previous 25 seconds. (**d–f**) t-SNE visualizations of each cell’s Ca^2+^ activity using features calculated using AQuA (*d*), ROIs (*e*), and tiles (*f*). k-means clustering boundary denoted as dashed line. (**g–h**) t-SNE plots using only subsets of AQuA-calculated features from (*a*) and (*d*). **(g)** t-SNE plot of only the features specific to AQuA and not shared with ROI or tile analysis. **(h)** Plot using only AQuA-extracted features that correspond to those in ROI- or tile-based analyses. **(i)** Comparison of difference between two clusters generated from the t-SNE analysis followed by k-means clustering. Note increased separation using AQuA-specific features compared to others. (One-tailed paired t-test, ***p<0.001)

**Supplemental Figure 9:**
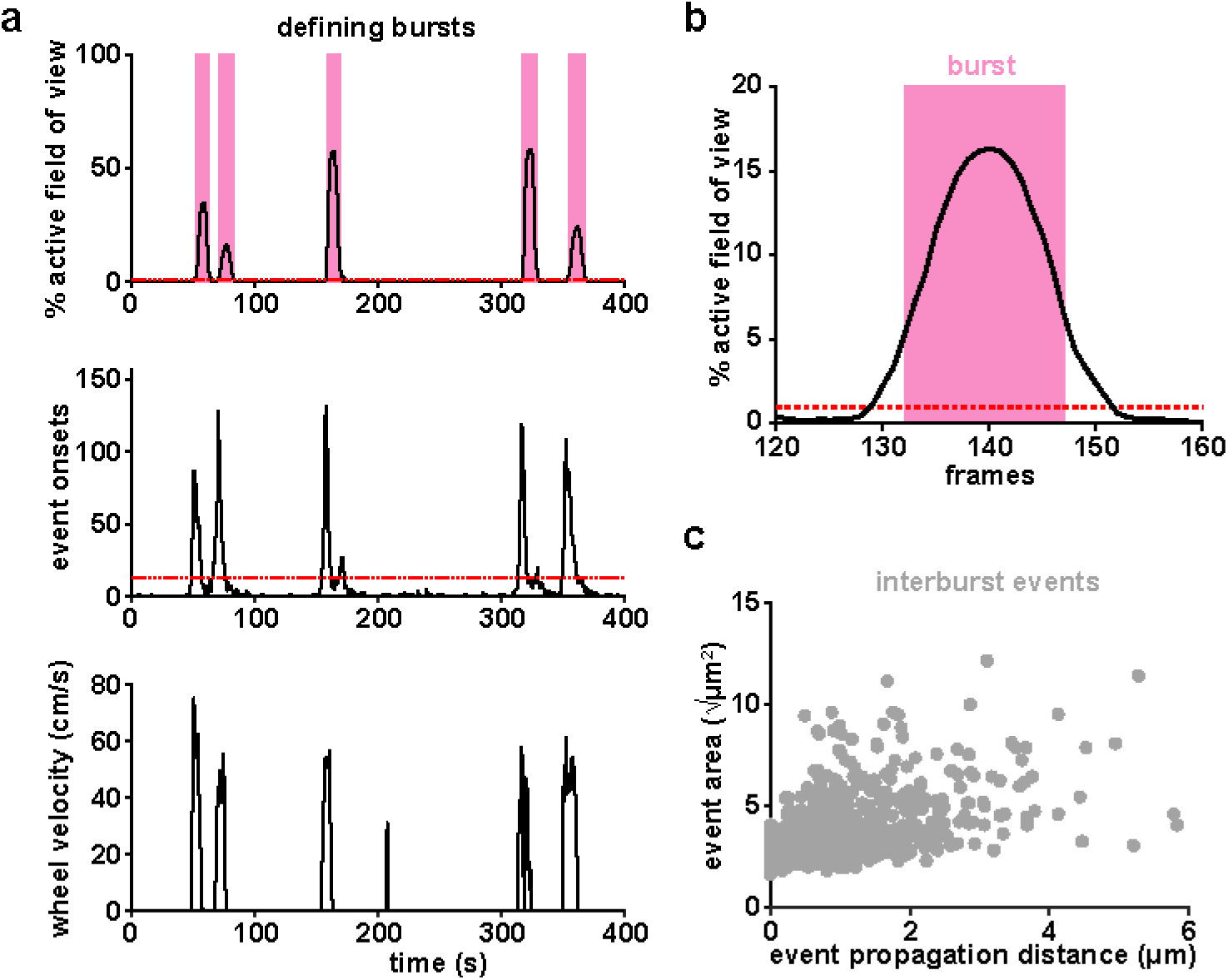
**(a)** Population Ca^2+^ events represented as two temporal traces: percentage of imaging field active (top) and number of AQuA event onsets (middle). Burst periods (pink) are defined from the top trace as periods when Ca^2+^ activity exceeds 1% of the active field of view (red dashed line, top), and exceeds 10% of the maximum number of event onsets (red dashed line, middle). Burst periods correlate with wheel velocity of the treadmill (bottom). **(b)** Burst onset is defined as the first frame in which 10% of the peak is exceeded and burst offset is defined as the last frame exceeding 10% of the peak. **(c)** The relationship between all interburst events’ total propagation distance and size, similar to the burst events plotted in Fig 4c.

**Supplemental Figure 10:**
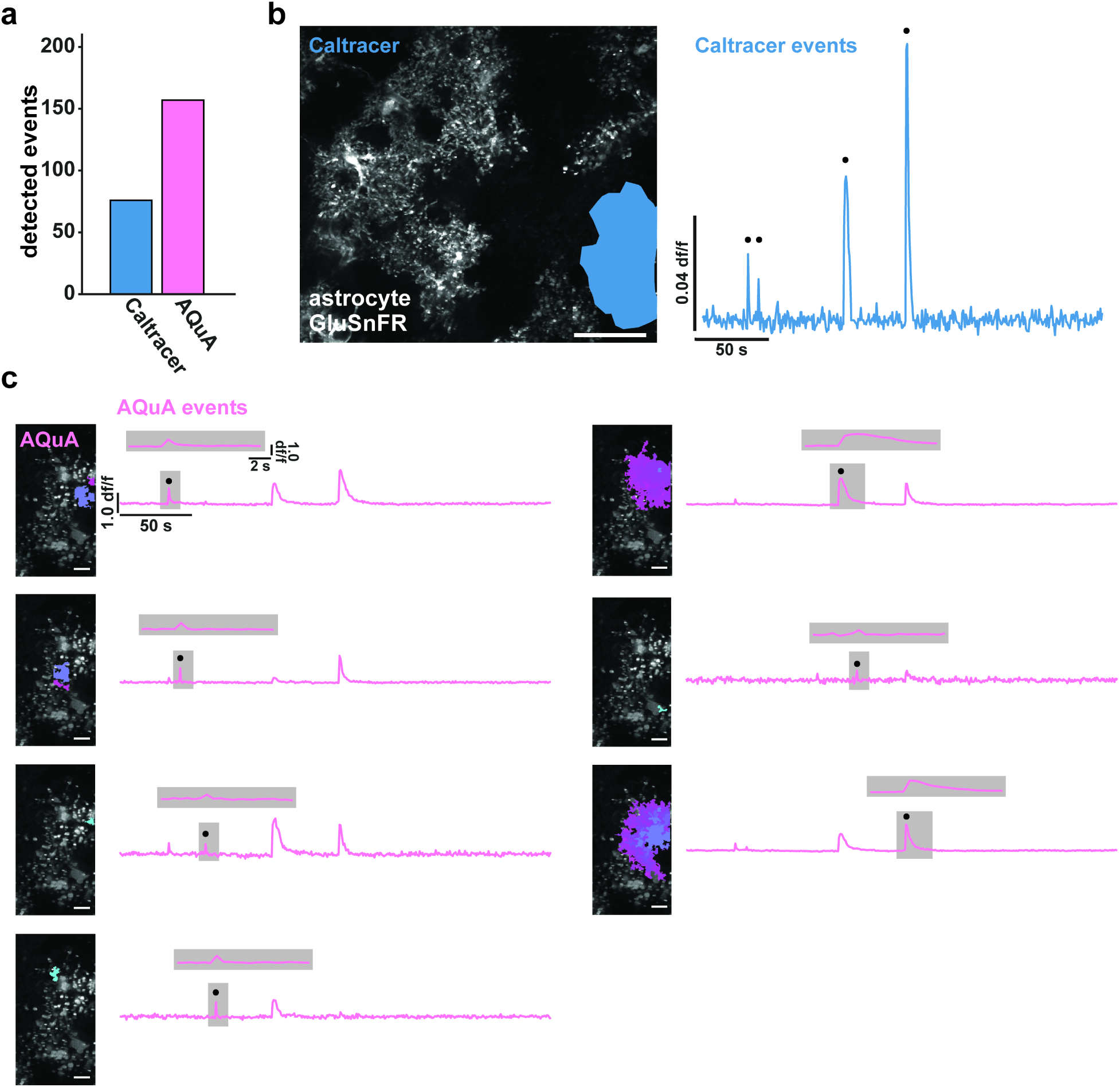
Comparison of AQuA and Caltracer for event detection of astrocytic GluSnFR signals. **(a)** Applied to the same data sets, AQuA detects 157 events, while Caltracer^2,9^, using a rising faces algorithm, detects 76 events with manually defined single-cell ROIs. **(b)** ROI example (left) and temporal trace with detected events marked by black circle using Caltracer software. Scale bar = 50μm. **(c)** AQuA-detected events from the same cell as in (b), and corresponding temporal traces (black dot, specific events shown above each trace). Scale bar = 10μm.

**Supplemental Figure 11:**
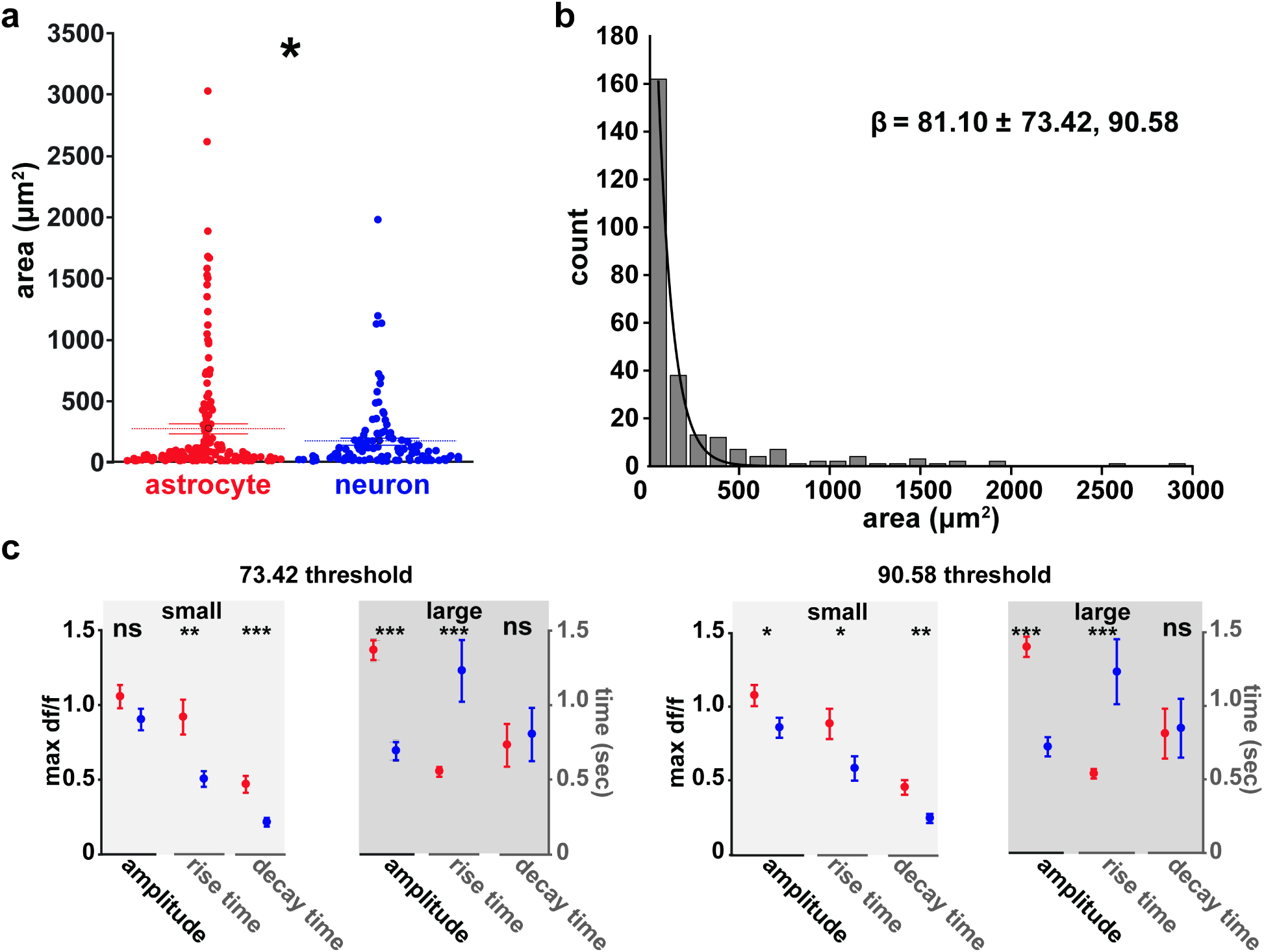
Differences in astrocytic and neuronal glutamate events are robust to varying size thresholds. Entire population (**a**) and distribution (**b**) of astrocytic and neuronal glutamate event size. Exponential fit and expected decay value shown with fit error. (**c**) Size thresholds set to lower (73.42, left) and upper (90.58, left) fit error bounds. Data are shown as mean ± SEM.

